# Efficient internalization of poly(benzyl malate) and poly(ethylene glycol)-*b*-poly(benzyl malate) copolymer based nanoparticles by human hepatic HepaRG cells and macrophages : Impact of nanoparticle functionalization by GBVA10-9 peptide on cell uptake

**DOI:** 10.1101/2024.07.26.605135

**Authors:** Nahas Hanadi, Saba Saad, Metlej Perla, Ribault Catherine, Vene Elise, Lepareur Nicolas, Cammas-Marion Sandrine, Loyer Pascal

**Author notes:** Corresponding authors: Sandrine Cammas-Marion and Pascal Loyer.

## Abstract

In the past years, we have designed biodegradable poly(benzyl malate) (PMLABe_73_) homopolymer and amphiphilic poly(ethylene glycol)-*b*-PMLABe (PEG_42_-*b*-PMLABe_73_) copolymer and several modified (co)polymers to produce biocompatible polymeric nanoparticles (NPs) capable of targeting hepatic cells *in vitro* with the goal to develop applications in the treatment of liver diseases. The current study aimed at comparing the uptake of PMLABe_73_ PEG_42_-*b*-PMLABe_73_-based NPs in human hepatic HepaRG cells, primary macrophages and peripheral blood mononuclear cells (PBMC). The uptake of NPs prepared from PEG_42_-*b*-PMLABe_73_ was significantly lower than that of PMLABe_73_ in both hepatic cells and macrophages. In addition, the NPs uptake by HepaRG cells was inversely correlated to the density of PEG present on their surface. In contrast, the internalization of with PMLABe-based NPs by human macrophages was not affected by low PEG densities, only uptake of fully pegylated PEG_42_-*b*-PMLABe_73_based-NPs was significantly decreased. Herein, we also showed that PMLABe_-_based NPs did not strongly accumulated in PBMC, T lymphocytes and neutrophils while monocytes showed slightly higher uptake of these NPs. Moreover, we further demonstrated that PMLABe-derived NPs by did not trigger inflammasome activation and secretion of pro-inflammatory cytokines neither in macrophages nor HepaRG cells. Then, we demonstrated that peptide GBVA10-9 derived from George Baker (GB) Virus A, known to exhibit a good hepatotropism did not significantly affect the uptake of PMLABe_73_-based NPs in HepaRG cells and macrophages, when grafted onto these NPs. The present results demonstrate that PMLABe-derived NPs are very efficiently internalized in both macrophages and hepatocytes but not in PBMC and reinforce our previous reports regarding their biocompatibility.

## I INTRODUCTION

Drugs administered by oral route and systemic injection distribute evenly throughout the body resulting in limited bioavailability deleterious, their rapid metabolism and elimination by the liver and kidneys and, in some cases in side-effects in healthy organs. Low bioavailability and inability to address chemotherapies to target tissues significantly contributes to the retrieval of promising molecules and low efficacy of approved drugs [Kola and Landis, 2004 ; Blanco et al., 2015]. In addition, the development of certain drugs is sometimes slowed down or even ended due to problems with their solubility or stability.

The field of nanotechnologies has grown exponentially in the past decades with the production of a plethora of nanovectors for multiple applications in medicine. The term of drug delivery nanovectors refers as to engineered molecular systems embedding pharmaceutical compounds in order to reduce their biotransformation and clearance while improving the therapeutic index by limiting the side effects of the bioactive molecules [Hoffman, 2008]. Drug delivery is thus a general concept that considers the interaction between the drug and transport system, the dosage and the route of administration. The development of efficient drug delivery systems requires the elaboration of biocompatible synthetic materials that self-assemble in aqueous solution to form nanoparticles (NPs) capable to encapsulate large amounts of bioactive compounds and to release of these pharmaceutical compounds in a controlled manner and in specific areas of the body [Torchilin, 2006 ; Hoffman, 2008]. In this context, NPs with a hydrophobic core are also an interesting alternative for the administration of lipophilic drugs that cannot be administered under a native form [Couvreur and Vauthier, 2006].

In the last two decades, an impressive literature reporting basic and translational research on NPs has led to multiple preclinical/clinical trials and the approval of several formulations by regulatory authorities for diagnosis and therapeutics in oncology [O’Brien et al., 2004 ; Stinchcombe, 2007 ; Anselmo and Mitragotri, 2021 ; Aldosari et al., 2021], gene therapies [Coelho et al., 2013 ; Kristen et al., 2019] and more recently the mRNA vaccines [Dolgin, 2021]. These breakthroughs result from optimized synthesis of macromolecules, improvement of their physicochemical features such as size, porosity, shape and surface properties that play an important role in drug encapsulation and delivery [Raemdonck et al., 2015 ; Topete et al., 2015 ; Stylianopoulos et al., 2015], and the formulation of NPs by microfluidics [Almeida et al., 2024]. Despite these successes, the use of drug delivery nanostructures in clinical protocols is far from being a generalized routine and developments of novel NPs are still required to overcome some biological barriers and limitations to address specific cellular targets such as solid tumors with poor prognosis [Blanco et al., 2015].

The first major limitation in NP-based therapies is the mononuclear phagocyte system (MPS), which refers to all immune cells with high phagocytic activity. Following systemic administration, NPs immediately undergo opsonization, the non-specific interactions with plasma proteins, and recognition by antibodies and proteins of the complement system to form the so-called “protein corona” coating all NPs injected *in vivo* or incubated with serum *in vitro* [Frank and Fries, 1991 ; Owens and Peppas, 2006 ; Tenzer et al., 2013]. This process is the first activation step of the innate immune system, which enhances the activity of the MPS to eliminate pathogens from the body. Although all factors controlling *in vivo* fate of NPs are probably not yet elucidated, NPs’ features have been optimized to reduce the opsonization and scavenging by MPS through the modulation of their size/shape [Blanco et al., 2015], surface charge [Arvizo et al., 2011], chemical structure [Mahon et al., 2012]. For instance, the addition of neutral polymers such as poly(ethylene glycol) (PEG) or dextran results in a “stealth” behavior towards opsonins, which reduces the activation of the complement [Coty et al., 2017], the MPS uptake and extends NP’s systemic lifetime [Tenzer et al., 2013 ; Kouser et al., 2018]. In addition, the PEG corona often increases the hydrodynamic diameter of the NPs thereby decreasing the renal clearance [Veronese and Pasut, 2005].

Another limiting biological barrier for site-specific targeting using NPs is the continuous endothelium of the blood vessels, which requires the NP’s translocation across endothelial cells to reach a given organ. The discovery of the Enhanced Permeability and Retention effect (EPR) [Maeda et al., 2013 ; Maeda et al., 2016] defined as the extravasation of macromolecules and NPs across the disorganized and/or fenestrated blood vessels within solid tumors, results in accumulation of NPs within the tumoral mass. Tumors also exhibit varying degrees of lymphatic drainage and macrophage infiltration which further increase accumulation of NPs in the vicinity of tumoral cells [Dai et al., 2018 ; Penn et al., 2018]. The rationale for drug delivery using NPs thus relies on this EPR-mediated passive targeting, [Blanco et al., 2015] and the efficient uptake of NPs by cancer cells [Sahay et al., 2010 ; Means et al., 2022]. While the EPR effect is well documented in murine models of cancer and widely accepted by the scientific community, the EPR effect in humans remains controversial, even though it is one of the conceptual pillars of the use NPs in oncology [Bertrand and Leroux, 2012 ; Youden et al., 2022]. In mouse models of xenografted tumors for which an EPR effect is attested, some studies have concluded that very low doses of injected NPs reached the tumor sites [Wilhelm et al., 2016 ; Dai et al., 2018; Penn et al., 2018]. In contrast, other publications using similar cancer models reported that the overall exposure of the tumor to NPs was ∼75% of the total amount of NPs injected in the blood [Price et al., 2020]. These contradictory conclusions highlighted the lack of knowledge about the pharmacokinetics and distribution of NPs, and questioned the relevance of the parameters for the evaluation of tumor targeting [McNeil, 2016].

In order to improve specific cell/tissue targeting, NPs have been functionalized with various protein ligands of membrane receptors differentially expressed in cells including short peptides [Dawidczyk et al., 2014 ; Gao et al., 2015 ; Shi et al., 2017 ; Zhu et al., 2018 ; Sun et al., 2018]. While some articles reported an increase in cell targeting mediated by peptide-decorated NPs, mainly in tumor models [Wicky et al., 2015 ; Zhu et al., 2018], others conclude to the low efficiency of the peptide functionalization because of the reduced diffusion of NPs into solid tumors and their internalization by macrophages within the tumors [Dai et al., 2018 ; Penn et al., 2018] and by the MPS especially in the liver and spleen [Ishida et al., 2006 ; Blanco at al., 2015].

After injection and opsonization, most NPs accumulate in the liver and spleen because of the liver sinusoids are highly specialized capillaries harboring large fenestrations in the endothelium and lacking basal lamina. This hepatic architecture greatly enhances the exchange between the liver parenchyma and the blood stream coming from the digestive tract and the hepatic artery [Jacobs et al., 2010] and favors the accumulation of NPs within the perisinusoidal space (space of Disse). In this small gap between the fenestrated endothelium and the trabecular hepatocytes, NPs are in close contact with hepatocytes, liver sinusoidal cells (LSECs), hepatic stellate cells (HSCs) and the liver resident macrophages or Küpffer cells, a major first line of the innate immunity. This active phagocytic activity in the normal liver [D’Addio et al., 2012 ; Bertrand et al., 2017] is a major “cell barrier” impairing long-term circulation of NPs and the use of nanotechnology-based therapy for targeting diseased organs. Phagocytosis by Küpffer cells is strongly correlated to the chemistry and surface charge, the size and shape of the NPs through the formation of the protein corona onto NPs and the fixation of antibodies and proteins of the complement [Gustafson et al., 2015]. In addition, some NPs activate the inflammasome, a major pathway of the innate immune system involved in the production of pro-inflammatory cytokines [Baron et al., 2015]. Conversely, authors have forecast that the use of NPs in other fields of clinical applications than oncology would be possibly translated to human liver diseases because of the “passive” accumulation of nanovectors in livers that do not undergo deep alterations of their architecture as observed in cancers [Reddy and Couvreur, 2011 ; Tacke, 2017 ; Zhang et al., 2016]. In this context, different studies have taken advantage of the active phagocytic activity of Küpffer cells to specifically deliver therapeutics to these cells infected with microorganisms such as bacteria, leishmaniasis and salmonellosis [Alving et al., 1978 ; Fattal et al., 1991], leading to strategies of immunomodulation using nanomedicine accumulating in the hepatic parenchyma [Pati et al., 2018 ; Luan and Ju, 2018]. It has also been demonstrated that hepatic accumulation of ultra-small superparamagnetic iron oxide particle (USPIO) is decreased in patients with Non-Alcoholic Steatohepatitis (NASH) and that USPIO-mediated magnetic resonance imaging can be used for diagnosis of NASH in human patients [Smits et al., 2016] further reinforcing the idea that liver homing of NPs is of great interest in human therapy beyond cancer treatment.

In the field of liver transplantation, the use of NPs has recently been evaluated mainly on rodent livers with promising results opening up interesting perspective for improving grafts [Yao and Martins, 2020]. Due to the increase in candidates for liver transplantation (LT), transplant teams had to broaden the acceptance criteria for so-called “expanded criteria” grafts [Nemes et al, 2016], which are more sensitive to ischemia-reperfusion injury generated during harvesting and the static preservation phase [Noack et al., 1993]. On a larger scale, perfusion of isolated organs using perfusion machines before revascularization during transplantation has demonstrated its effectiveness in reducing the creation of ischemia-reperfusion lesions resulting in an improvement in the recovery of graft function, the incidence of complications and an improvement in its survival [Schlegel et al., 2013 ; van Rijn et al., 2021 ; Dutkowski et al., 2015]. To date, only one pilot study evaluating the use of nanovectors on human livers refused for transplantation has been carried out [Del Turco et al., 2022]. This study used non-degradable cerium oxide NPs conjugated to albumin, administered using a homemade perfusion machine model. The interest of these NPs is the presence of Ce^3+^/Ce^4+^ ions on their surface producing anti-oxidant and anti-inflammatory effects, which persist over time due to the very high stability of these structures. In this study, the authors demonstrate the internalization of these NPs in liver cells, including hepatocytes. These NPs improved the redox status with a maintenance of the glutathione pool and an increase in catalase activity but without a positive effect on the production of pro-inflammatory cytokines. We postulate that the administration of biocompatible NPs, specifically targeting hepatocytes, cholangiocytes and hepatic macrophages coupled with the use of perfusion machines could allow innovative targeted therapies and regenerative medicine for damaged liver grafts and ultimately to increase the number of transplantation procedures, to improve the quality and the resumption of functions of the grafts and consequently the performance of liver transplantation.

The Amphiphilic block copolymers are promising compounds for drug delivery since they form NPs or micelles in aqueous solutions with a hydrophobic inner-core surrounded by a hydrophilic corona. Our laboratory has developed poly(malic acid) (PMLA) derivatives for liver targeting drug delivery systems [Cammas et al., 2000 ; Huang et al., 2012 ; Loyer and Cammas-Marion, 2014 ; Casajus et al., 2018]. Amphiphilic derivatives of PMLA constituted by a PEG hydrophilic block and a poly(benzyl malate) (PMLABe) hydrophobic segment self-assemble to form PEG_42_-*b*-PMLABe_73_ micelles that show very low cytotoxicity levels towards hepatic cells and macrophages [Huang 2012 ; Casajus et al., 2018]. In previous reports, we also grafted Circumsporozoite protein of *Plasmodium berghei*-(CPB) and George Baker Virus A-10-9-(GBVA10-9) derived peptides, which showed a good hepatotropism towards hepatic cells [Brossard et al., 2021 ; Vène et al., 2022, Brossard et al., 2022].

In this report, our first objective was to better characterize the uptake of poly(benzyl malate) and poly(ethylene glycol)-*b*-poly(benzyl malate) copolymer based NPs by human hepatic cells and macrophages and to study for the first time the internalization of these NPs by peripheral blood mononuclear cells. We also determined the impact of the NP’s functionalization by the GBVA10-9 peptide. The second objective was to set up a coculture system combining hepatic HepaRG cells and human macrophages in order to determine whether the peptide-functionalization of PMLABe and PEG-*b*-PMLABe-based NPs with GBVA10-9 peptide could favor the uptake by hepatic cells over that of macrophages in this *in vitro* model allowing cell competition for the internalization of GBVA10-9-decorated NPs.

## II MATERIALS AND METHODS

### 2.1 Materials

#### Dynamic Light Scattering (DLS)

DLS measurements were performed on a Nano-sizer ZS90 (Malvern, Worcestershire, UK) at 25°C, with a He-Ne laser at 633 nm and a detection angle of 90°C. Three runs of 70 scans each were performed on each NPs suspension, and average values of hydrodynamic diameter (Dh) and dispersity (PDI) were given. The size distribution reports were given by Intensity.

#### Size Exclusion Chromatography (SEC)

Weight average molar mass (Mw) and dispersity (Đ = Mw/Mn) values were measured by SEC in THF at 40 °C (flow rate = 1.0 mL/min) on a GPC2502 Viscotek apparatus equipped with a refractive index detector Viscotek VE 3580 RI, a guard column Viscotek TGuard, Org 10 x 4.6 mm, a LT5000L gel column 300 x 7.8 mm and a GPC/SEC OmniSEC Software (Malvern, Worcestershire, UK). The polymer samples were dissolved in THF (2 mg/mL). All elution curves were calibrated with polystyrene standards.

#### Differential Scanning Calorimetry (DSC)

Glass transition temperature (Tg) of the (co)polymers was measured by DSC. Measurements were acquired on a DSC Q2000 apparatus from TA Instruments under nitrogen flow at heating rate 10°C/min from -80 to 180 °C.

#### Flow cytometry and microscopy

The cell uptake of fluorescent NPs labelled with the lipophilic fluorescent dye 1,1′-dioctadecyl-3,3,3′,3′-tetramethylindodicarbocyanine perchlorate (DiD_Oil; Thermofisher Scientitic, Wavelength: excitation 644 nm; emission 665 nm, ε = 236.000) was quantified by flow cytometry using 2 different analyzers from the cytometry core facility of the Biology and Health Federative research structure Biosit (University of Rennes, France): FACSCalibur and LSRFortessa™ X-20 cytometers (Becton Dickinson, Becton Drive Lake, NJ, USA) and data were analyzed using CellQuest and FACSDiva^TM^ softwares, for these two appartus, respectively (Becton Dikinson). Fluorescent cells were visualized using a Zeiss AxioVert A.1 microscope coupled with a Colibri.2 illumination system (Carl Zeiss Microscopy GmbH, Germany).

#### ELISA assays

For ELISA assays, the optical absorbance was measured on a microplate reader Multiskan FC (ThermoScientific).

#### Western blotting

Electrophoresis and protein transfers were performed on XCell SureLock^TM^ and iBlot2^®^ apparatus (Life Technologies). Acquisitions of gels stained with coomassie blue and immunoblotting detection by chemioluminescence were performed using VisionCapt and Chemi-Smart 5000 systems (Vilber Lourmat), respectively.

### 2.2 Reagents

All chemicals were used as received. α-maleimide,ω-carboxylic acid PEG_62_ (Mal-PEG_62_-COOH, Mw = 3,000 g/mol, n = 62) and α-methoxy,ω-carboxylic acid PEG_42_ (MeOPEG_42_-COOH, Mw = 2,015 g/mol, n = 42) were purchased from Iris Biotech GmbH (Marktredwitz, Germany). Tetraethylammonium benzoate, tetraethylammonium hydroxide and 6-maleimidohexanoic acid (Mal-Hex-COOH) were purchased from Sigma-Aldrich (Saint-Louis, Mo, USA). Peptides were provided by Eurogentec (Liege, Belgium). 1,1’-Dioctadecyl-3,3,3’,3’-tetramethylindodicarbocyanine perchlorate (DiD Oil) was purchased from Invitrogen (Thermo Fisher Scientific, Illkirch Graffenstaden, France). Solvents were purchased from Sigma-Aldrich (Saint Quentin Fallavier, France).

Phosphate-buffered saline (PBS), William’s E medium, RPMI 1640, penicillin–streptomycin, L-glutamine and trypsin were purchased from ThermoFisher Scientific (Illkirch Graffenstaden, France). Fetal calf serum (FCS) FetalClone III^®^ and BioWhittaker^®^ were from Hyclone (Logan, UR, USA) and Lonza (Verviers, Belgium), respectively. Hydrocortisone hemisuccinate was from Serb (Paris, France). MOPS-SDS buffer was purchased from Amresco (OH, USA). Tris-buffered saline (TBS) was from GE Healthcare (Aulnay Sous Bois, France). Bovine serum albumin was from Eurobio (Les Ulis, France). Ultrapure Escherichia coli O111:B4 LPS was purchased from InvivoGen (Toulouse, France) and recombinant human granulocyte macrophage colony-stimulating factor (rhGM-CSF) from R&D Systems Europe (Lille, France). Insulin were obtained from Sigma-Aldrich (Saint Louis, MO, USA). Carboxylate-modified fluorescent (yellow-green) FluoSpheres^®^ (50 and 100 nm) and 1,1′-dioctadecyl-3,3,3′,3′-tetramethylindodicarbocyanine perchlorate (DiD-Oil) were purchased from Molecular Probes (Eugene, OR, USA). Monosodium urate (MSU) crystals were prepared by recrystallization from uric acid, as previously described [Gicquel et al., 2015]. Goat antiserum to human albumin (1140V7) was from Kent Laboratories (Redmond, WA, USA), rabbit anti-complement C3 (sc-31300) and anti-Apolipoprotein (B-10) were from Santa Cruz (distributed by CliniSciences, Nanterre, France) and anti-human immunoglobulins was purchased from Amersham (RPN 1003). Horse radish peroxidase labeled secondary antibodies were from Dako (Denmark).

### 2.3 Formulation of PMLABe_73_ derivatives-based nanoparticles

PMLABe_73_, Mal-PMLABe_73_, PEG_42_-*b*-PMLABe_73_ and Mal-PEG_62_-*b*-PMLABe_73_ were first synthesized by anionic ring opening polymerization (aROP) of benzyl malolactonate (MLABe) using tetraethylammonium benzoate, tetraethylammonium maleimidohexanoate, tetraethylammonium salt of ω-methoxy,α-carboxylate-PEG_42_ and tetraethylammonium salt of α-maleimide,ω-carboxylic acid PEG62as initiator, respectively, following the procedure previously described [Brossard et al., 2021; Brossard et al. 2022] and summarized in Supplementary Information 1A. All the synthesized (co)polymers were characterized by ^1^H NMR (structure and molar mass), SEC (Mw and Ð) and DSC (Tg) as already reported [Brossard et al., 2021; Brossard et al. 2022].

Nanoparticles (NPs) encapsulating the fluorescence probe DiD Oil were then prepared using the nanoprecipitation technique [Brossard et al., 2021; Brossard et al. 2022], adapted from the method described previously [Thioune et al., 1997]. Generally, 5 mg of (co)polymers [100wt% PMLABe_73_ (PMLABe_73_-NPs), 100wt% MeOPEG_42_-*b*-PMLABe_73_ (PEG_42_-*b*-PMLABe_73_-NPs), 75wt% PMLABe_73_ + 25wt% MeOPEG_42_-*b*-PMLABe_73_ (PMLABe_73_/PEG 75/25-NPs), 50wt% PMLABe_73_ + 50wt% MeOPEG_42_-*b*-PMLABe_73_ (PMLABe_73_/PEG 50/50-NPs), 90wt% PMLABe_73_ + 10wt% Mal-PMLABe_73_ (PMLABe_73_/Mal-PMLABe_73_ 90/10-NPs), or 90wt% PMLABe_73_ + 10wt% Mal-PEG_62_-*b*-PMLABe_73_ (PMLABe_73_/Mal-PEG_62_-*b*-PMLABe_73_ 90/10-NPs)] were solubilized in DMF followed by the addition of a given volume of the DiD Oil stock solution at a concentration of 0.1 mg/mL (the amount of DiD Oil representing 0.1 wt% of the (co)polymers’ mass, i.e. 0.05 mg). The final volume of DMF was always 150 µL. This solution was rapidly added in ultra-pure water (1 or 2 mL) under vigorous stirring. After 10 min of stirring at room temperature, the DiD Oil-loaded NPs suspension were passed through a Sephadex G25 leading to the obtaining of 3.5 mL of DiD Oil-loaded NPs suspensions [Brossard et al., 2021; Brossard et al. 2022]. The obtained NPs suspensions were characterized by DLS (Table 1).

**Table 1:** Characteristics of the PMLABe-derived nanoparticles. . Nanoparticle (NP) composition indicating the (co)polymer(s) used to formulate NPs: PMLABe_73_, PEG_42_-*b*-PMLABe_73_ and their maleimide-modified derivates used in various proportions to prepare corresponding NPs. Dh: Hydrodynamic diameter measured by DLS. PDI: Polydispersity index of the NP size measured by DLS.

| **NPs composition** | **Dh (nm)** | **PDI** |
| --- | --- | --- |
| PMLABe<sub>73</sub>[DiD Oil] | 120 | 0.17 |
| PEG<sub>42</sub>-*b*-PMLABe<sub>73</sub>[DiD Oil] | 106 | 0.19 |
| PEG<sub>42</sub>-*b*-PMLABe<sub>73</sub>/PMLABe<sub>73</sub>[DiD Oil] 25/75 | 119 | 0.20 |
| PEG<sub>42</sub>-*b*-PMLABe<sub>73</sub>/PMLABe<sub>73</sub>[DiD Oil] 50/50 | 115 | 0.16 |
| PMLABe<sub>73</sub>/ GBVA10-9-PMLABe<sub>73</sub>[DiD Oil] 90/10 | 139 | 0.42 |
| PMLABe<sub>73</sub>/ GBVA10-9-PEG<sub>62</sub>-*b*-PMLABe<sub>73</sub>[DiD Oil] 90/10 | 142 | 0.38 |

The GBVA10-9 C-terminated with a thiol group (GBVA10-9-SH) in solution in PBS was added to the maleimide-decorated NPs suspensions (PMLABe_73_/Mal-PMLABe_73_ 90/10-NPs, and PMLABe_73_/Mal-PEG_62_-*b*-PMLABe_73_ 90/10-NPs) through the Michael addition using conditions described previously [Brossard et al. 2022], thus leading to the GBVA10-9-decorated NPs which were characterized by DLS (Table 1).

### 2.4 Opsonization of PMLABe, PEG-b-PMLABe based NPs

The opsonization of NPs was studied using a protein adsorption assay followed by western blot analysis. PMLABE and PEG_62_-*b*-PMLABe_73_–based NPs (functionalized or not) were incubated in William’s E medium supplemented with 10% human serum during 24h at 37°C under 5% CO_2_ humidified atmosphere. NPs were collected by centrifugation at 14,000 g for 30 min while agarose beads were spun down at 5,000 g for 1 min, at 4°C. The pellets were washed once with cold PBS (500µL) prior to denaturation of micelles and bound proteins with 50 µL of denaturating buffer (Tris-HCl 100 mM, pH 6.8, bromophenol blue 0.2%, sodium dodecyl sulfate 8%, glycerol 20%, and β-mercaptoethanol 5%) and 50µL of MOPS/SDS buffer pH 7.7 (Thermofisher Scientific, USA). Samples were boiled in water bath for 10 min. Standard PageRuler^TM^ Plus prestained protein ladder (Thermofisher Scientific) and protein samples were loaded and separated by electrophoresis on polyacrylamide gels (iD PAGE gel, 4-12%, Eurogentec, Belgium) and then transferred to nitrocellulose membranes (iBlot® 2NC Mini Stacks, Thermofisher Scientific, USA). The membranes were blocked with 3% bovine serum albumin (BSA) Fraction V (Eurobio) in 1X Tris-buffered saline, 0.1 % Tween 20 (TBST) at room temperature (RT) for 1 h, then incubated overnight at 4 °C with the following primary antibodies diluted in TBST containing 3% BSA: mouse anti-human complement C3 proteins (*B-9)*, mouse anti-human apolipoprotein A-I (B-10), goat anti-human albumin (1140V7) and goat anti-human IgG (A-0293). Following washes three times with TBST, the membranes were then incubated for 1 h at RT with appropriate horseradish peroxidase (HRP) secondary antibodies: polyclonal rabbit anti-mouse and polyclonal goat anti-rabbit. After incubation, the membranes were washed three times with TBST and developed using SuperSignal^TM^ WestDura chemiluminescent substrate kit for HRP detection according to the manufacturer’s instructions. The proteins were visualized with the Fusion FX system (Vilber-Lourmat, Germany).

### 2.5 Cell culture and cell uptake of NPs

HepaRG cells were cultured as previously described [Corlu and Loyer, 2015]. Briefly, progenitors HepaRG were cultured in William’s E medium supplemented with 2 mM L-glutamine, 50 IU/mL penicillin, 50 µg/mL streptomycin, 5 mg/L insulin, 10^-5^M hydrocortisone hemisuccinate and 10% FBS. To obtain differentiated HepaRG cells, progenitors were cultured during 14 days to obtain confluent quiescent cells and maintained for 2 additional weeks in medium supplemented with 2% DMSO.

Human peripheral blood mononuclear cells (PBMC) were isolated from buffy coat of healthy donors (Etablissement Français du Sang, Rennes, France) by centrifugation on UNI-SEP maxi U10 (Novamed). Monocytes (CD14^+^) were isolated using anti-human CD14 antibodies conjugated magnetic MicroBeads (Miltenyi Biotec SAS, Paris, France) and were plated at a density of 0.5×10^5^ cells per well in 48-well plates. Human macrophages were obtained after differentiation of monocytes with 50 ng/mL rhGM-CSF in RPMI 1640 medium supplemented with 5 IU/mL penicillin and 5 mg/mL streptomycin, 2 mM L-glutamine and 10% FBS during 7 days, as previously described [Vène et al., 2016 ; Vène et al., 2022]. After 7 days of differentiation, the culture medium was removed and 1×10^5^ cells HepaRG cells expressing the Green Fluorescent Protein (GFP) were added in human macrophage wells. RPMI 1640 medium supplemented with 10% FBS, 50 IU/mL penicillin, 50 μg/mL streptomycin, and 2 mM L-glutamine was used for the coculture. GFP-expressing HepaRG cells were produced by lentiviral transduction of proliferating cells plated at low cell density (10^5^ cells per well in 24-well plates) with pre-made lentiviral particles (ILV-EF1-GFP) obtained from Flash Therapeutics (Toulouse, France). All cell types were incubated at 37°C with 5% humidified CO_2_.

### 2.6 Cell uptake of PMLABe-based NPs and FluoSpheres^®^

Control and peptide-functionalized fluorescent NPs were prepared as described in section 2.3. Culture medium was withdrawn and replaced by 500 µL (in 24-well plates) of fresh culture media containing NPs at a final concentration of 25 μg/mL in copolymer corresponding to 5.10^10^ NPs/mL.

After incubation at 37°C in a humidified atmosphere of 5% CO_2_ for various time points, cell monolayers were washed twice with PBS and photographs were acquired using fluorescence microscope. The cells were detached with trypsin-EDTA and resuspended in complete medium for flow cytometry analysis. Dot plots of forward scatter (FSC: x axis) and side scatter (SSC: y axis) allowed to gate the viable single cells (Supporting information 2). Untreated cells were used to determine autofluorescence, arbitrary set at ∼30 for all cell types. The fluorescence emitted by NPs encapsulating DiD Oil was detected using the APC-A channel. For GFP expressing HepaRG cells, the GFP was detected on FITC channel to gate hepatic cells and to discriminate human macrophages. The percentage of positive cells and the mean of fluorescence was expressed as fluorescence intensity of the single cell population (Supporting information 2). The effects of peptides on cell uptake of peptide-decorated NPs was evaluated with the percentage of positive cells and the mean of fluorescence for cells incubated with peptide functionalized streptavidin compared to those of NPs without peptides. For the cell uptake assay of FluoSpheres^®^, human macrophages and HepaRG cells were incubated with FluoSpheres^®^ according to the manufacturer instructions.

To evaluate the influence of the opsonization on cellular uptake, the assay described above was modified by incubating DiDoil-loaded NPs or FluoSpheres^®^ in culture media without fetal calf serum. The fluorescence emitted by the cells was analyzed by flow cytometry.

### 2.7 Detection of CD3+ lymphocytes and CD66+ neutrophils by plow cytometry

Cell number was determined, and 10^6^ cells were centrifuged at 300g for 10 minutes. Supernatant was discarded and cells were resuspended in 98 μl of phosphate-buffered saline (PBS) pH 7.2, 0.5% bovine serum albumin (BSA), and 2 mM EDTA. Two μl of CD3 antibody, anti-human, FITC, REAfinity™ (Miltenyi Biotech, reference 130-113-138) or CD66abce antibody, anti-human, PE, REAfinity™ (Miltenyi Biotech, reference 130-124-512) were added. REA Control antibody, human IgG1, FITC, REAfinity™ (Miltenyi Biotech, reference 130-113-437) and REA Control antibody, human IgG1, Vio_®_ Bright B515, REAfinity™ (Miltenyi Biotech, reference 130-113-445) were used as negative control for CD3 and CD66abce, respectively. Cells and antibody mix were then incubated for 10 minutes in the dark at 4°C. Cells were washed with 1-2 mL of buffer and centrifuged at 300xg for 10 minutes. Supernatant was discarded and cells were fixed with formaldehyde 4% prior to flow cytometry analysis.

### 2.8 Quantification of cytokines by ELISA assay

Macrophages and HepaRG cells were incubated during 24h with 0.1 µg/mL ultrapure lipopolysaccharide (LPS) for inflammation priming. Then, the culture media were discarded and cells were treated overnight with NPs, FluoSpheres^®^ or MSU 250 µg/mL. Production of cytokines was evaluated by quantification of interleukin-1β (IL-1β), IL-1α and IL-6 levels in culture supernatants of primed cells and cells incubated with NPs but without LPS treatments using Duoset ELISA kits, according to the manufacturer’s instructions.

### 2.9-Statistical analyses

Quantitative data were expressed as mean ± standard deviation (SD). Statistical analyses were performed using GraphPad Prism version 5.0 (GraphPad Software, USA). Differences between two groups were analyzed using two-tailed Mann-Whitney *U* test. A non-parametric Kruskal-Wallis test with Dunns’ post-test was used to compare means of more than two groups. Significant differences are presented as * p<0.05, ** p<0.01, *** p<0.001, otherwise: not significant.

## III RESULTS

### 3.1 In vitro cell uptake of PMLABe_73_ and PEG_42_-b-PMLABe_73_-based NPs by HepaRG cells and macrophages

The homopolymer PMLABe_73_ and block copolymer PEG_42_-*b*-PMLABe_73_ were synthesized by anionic ring opening polymerization (aROP) of benzyl malolactonate (MLABe) as described previously [Brossard et al., 2021; Brossard et al. 2022] and summarized in **Supporting Information 1A**. In a first step, PMLABe_73_- and PEG_42_-*b*-PMLABe_73_-based NPs (100% of either PMLABe_73_- or PEG_42_-*b*-PMLABe_73_-derived NPs) encapsulating the fluorescence probe DiD Oil were prepared using the nanoprecipitation technique. The homopolymer PMLABe_73_ and amphiphilic block copolymers PEG_42_-*b*-PMLABe_73_ self-assembled in aqueous solutions to form well-defined macromolecular NPs with hydrodynamic diameters of 120 and 106 nm, respectively (**Table 1**) while polydispersity indexes of ∼ 0.15 to 0.2 evidenced homogenous micelle formulations, in agreement with NPs obtained in our previous reports [Huang et al., 2012 ; Casajus et al., 2018 ; Brossard et al., 2021; Brossard et al. 2022].

The incubation of the NPs with human macrophages and HepaRG cells was performed overnight and cell uptake of PMLABe_73_- and PEG_42_-*b*-PMLABe_73_-based NPs was measured by the detection of the DiD oil encapsulated into the NPs using flow cytometry and fluorescence microscopy (**Figure 1**). The percentages of DiD oil positive cells, which had internalized PMLABe_73_ and PEG_42_-*b*-PMLABe_73_ -based NPs, and the means of fluorescence of cell populations were obtained from the flow cytometry data and represented in histograms and chart, respectively (**Figure 2**). The cell uptake of PMLABe_73_- and PEG_42_-*b*-PMLABe_73_-based NPs in primary human macrophages and HepaRG cells was compared to that of green carboxylate-modified polystyrene microspheres (FluoSpheres®) of 20 and 100nm (**Figures 1 and 2**).

**Figure 1:**
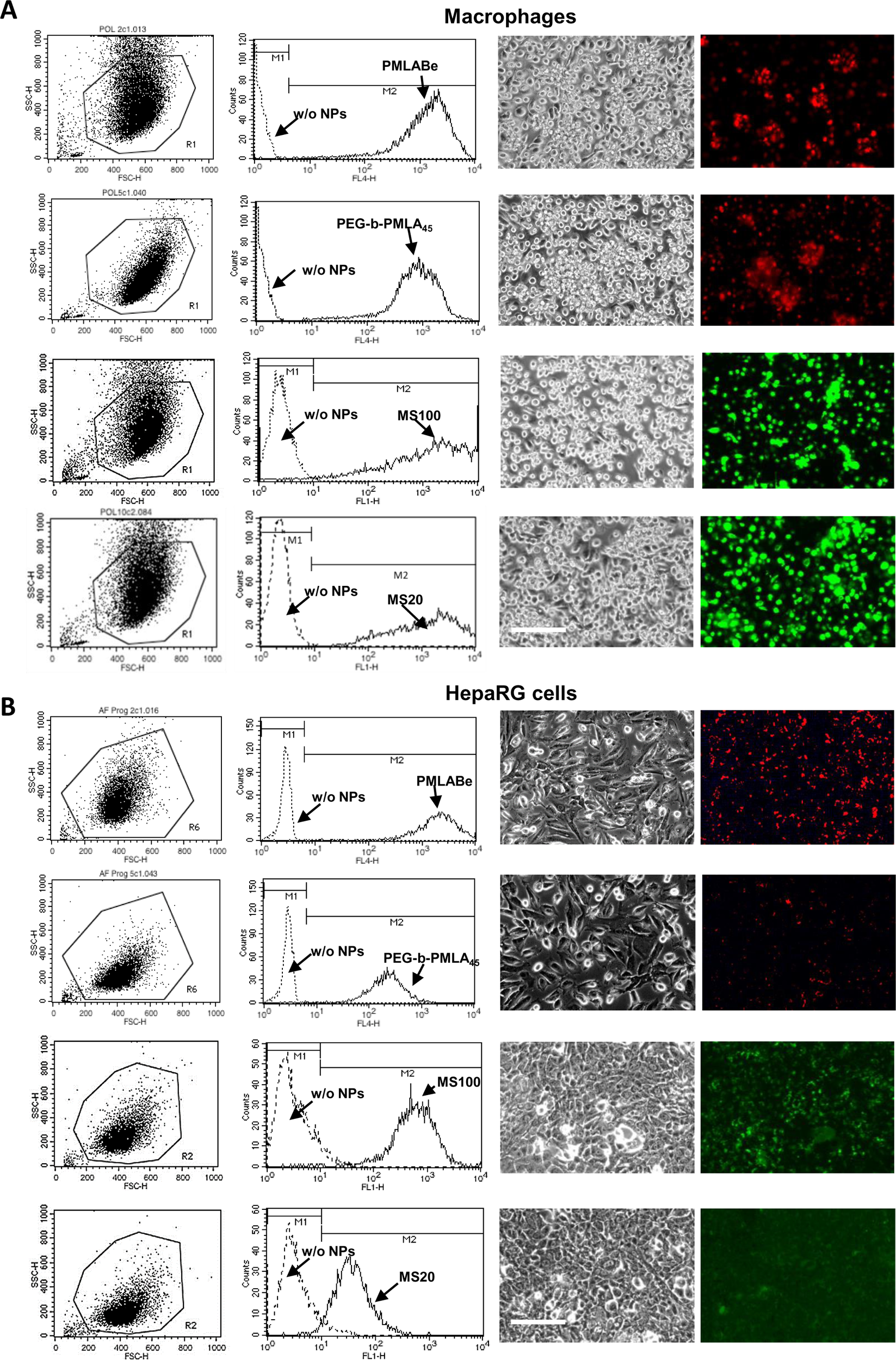
Uptake of PMLABe_73_-, PEG_42_-b-PMLABe_73_-based NPs and FluoSpheres^®^ in primary macrophages and HepaRG cells by flow cytometry and fluorescence microcopy. The flow cytometry analysis was performed using 10^4^ gated viable cells (gate R1 on the side scatter versus forward scatter dot plots, left column). The intrinsic FL4-H fluorescence of macrophages and HepaRG cells was set up using cells that were not incubated with NPs (w/o NPs, dotted line histograms) to define the M1 gate corresponding to negative cells. The fluorescence of macrophages and HepaRG cells incubated with DiD oil loaded NPs prepared from PMLABe_73_-homopolymer and PEG_42_-b-PMLABe_73_ copolymer and fluorescein-labelled FluoSpheres^®^, was measured using the FL4-H and FL1-H channels, respectively, to quantify the positive cells in the M2 gate. Only overlay histograms of cells w/o NPs and cells incubated with PMLABe-based NPs and FluoSpheres^®^ beads of 20 nm (MS20) and 100 nm (MS100) are presented. Flow cytometry was performed using a FACSCalibur analyzer (Becton Dickinson). Fluorescence DiD oil loaded NPs (red) and FluoSpheres^®^ microspheres (green) in macrophages and HepaRG cells were also detected by fluorescence microscopy: live cells in phase contrast microscopy are presented in the third column and the corresponding fluorescence photographs resulting from the accumulation of NPs into the cells are presented in the fourth column (magnification bar: 100μm).

**Figure 2:**
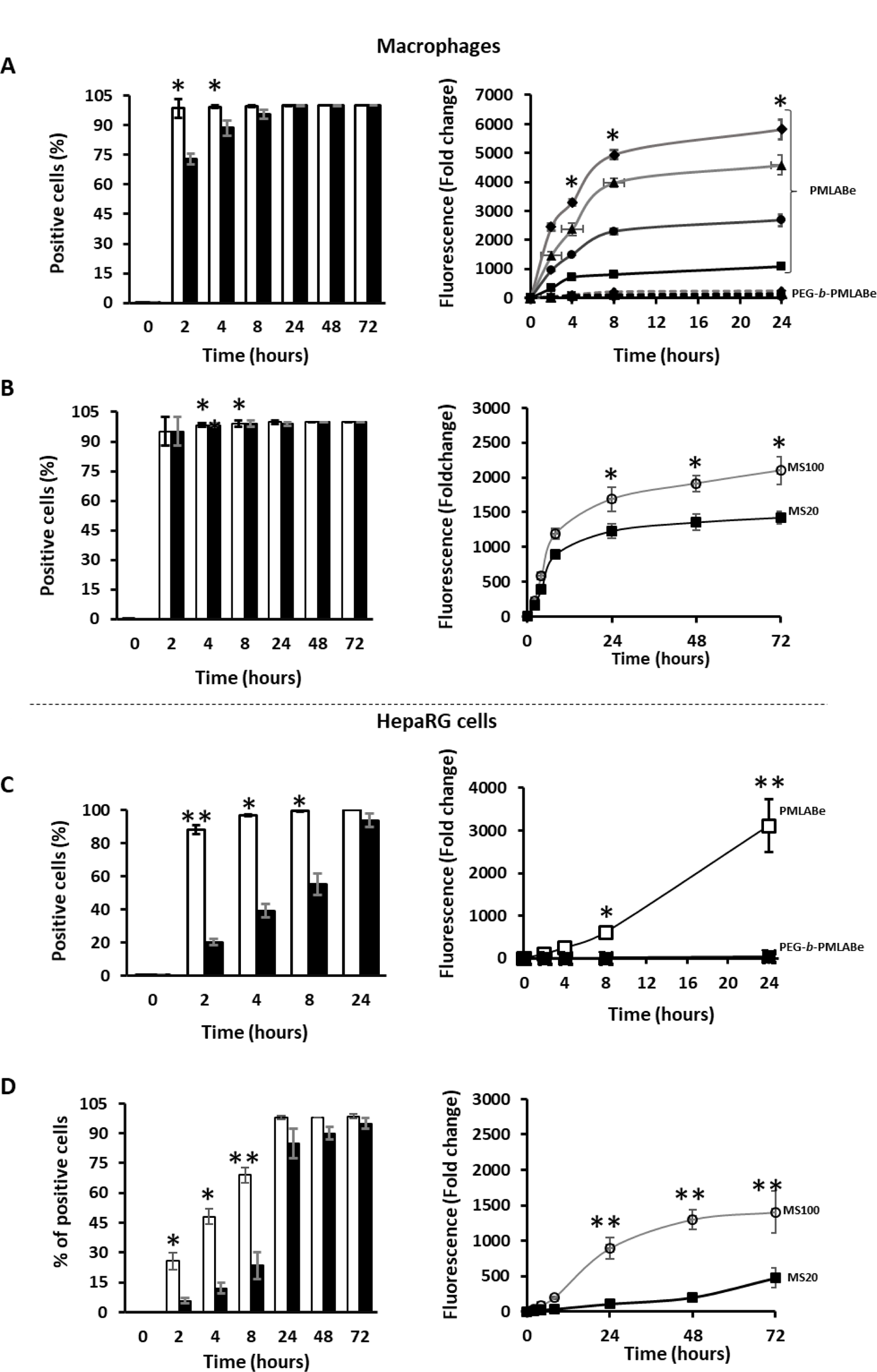
Quantification of the uptake of PMLABe_73_-, PEG_42_-b-PMLABe_73_-based NPs and FluoSpheres^®^ in primary macrophages and HepaRG cells by flow cytometry. The histograms represent the percentage of positive macrophages (A, B) and HepaRG cells (C, D) detected in the M2 gate (see Figure 1) after incubation for various times with PMLABe_73_-, PEG_42_-b-PMLABe_73_-based NPs (A, C) and FluoSpheres^®^ (MS100 or 20nm NPs). The curves represent the mean of fluorescence expressed in fold change of the background fluorescence measured in non-incubated cells. As control, cells were not incubated with NPs and their background fluorescence was the mean of the all cell populations. Macrophages prepared from four healthy donners were used to measure the uptake in 3 to 6 independent culture wells. Three independent cultures of HepaRG were performed to measure the uptake in 6 to 9 independent culture wells. * *p* < 0.05, ** *p* < 0.01.

As demonstrated by the flow cytometry histograms and fluorescence microscopy, a very efficient uptake of PMLABe_73_-, PEG_42_-*b*-PMLABe_73_-based NPs and FluoSpheres® by macrophages and HepaRG cells was observed after an overnight incubation of NPs with the tow cell types (**Figure 1**). Nearly all macrophages and HepaRG cells were positive following incubation with PMLABe_73_-, PEG_42_-*b*-PMLABe_73_-based NPs and FluoSpheres®. The time course study of the NP’s uptake showed significant differences between the uptake in macrophages and HepaRG cells and between the PMLABe_73_-, PEG_42_-*b*-PMLABe_73_-based NPs and FluoSpheres® (**Figure 2**). The uptake of PMLABe_73_-based NPs is much greater than that of PEG_42_-*b*-PMLABe_73_-based NPs for the two cell models, indicating that the presence of a PEG corona on the surface of these NPs strongly reduced their internalization. Our data also evidenced quantitative variations in the uptake of PMLABe_73_-based NPs between the different macrophage cultures prepared from four independent healthy donors visualized from important differences in fluorescence intensities (**Figure 2A**), which demonstrated variable interindividual NP’s internalization capacities among these donors. In contrast, the uptake of FluoSpheres® by the same four donors showed quite similar fluorescence intensities allowing to combine the values of the donors and resulting in narrow standard deviations (**Figure 2B**).

The time-course study showed that macrophages internalized very actively PMLABe_73_-, PEG_42_-*b*-PMLABe_73_-based NPs and FluoSpheres® in the first hours of incubation since nearly 95% of cells were positive at 8h (**Figure 2A**). Then, at 24h, while nearly all macrophages had internalized PMLABe_73_-, PEG_42_-*b*-PMLABe_73_-based NPs, the fluorescence intensities reflecting the accumulation of NPs weakly increased (**Figure 2A**). Similarly, the uptake of FluoSpheres® increased in a much lower extend compared to the fast internalization during the first 8h of incubation (**Figure 2B**). In addition, the means of fluorescence were significantly different in macrophages incubated with these microspheres of 100 and 20 nm In hepatic HepaRG cells, the internalization of PMLABe_73_-, PEG_42_-*b*-PMLABe_73_-based NPs and in a lesser extend for FluoSpheres® was more linear with the time of incubation (**Figure 2C**). The uptake of PMLABe_73_-based NPs was much higher both on percentages of positive cells and fluorescence intensities than the internalization of PEG_42_-*b*-PMLABe_73_-derived NPs. At 8h, all HepaRG cells were positive for the DiD-Oil labelling but their mean of fluorescence intensities was weaker than that found in macrophages. While 60 to 80% of macrophages had internalized PEG_42_-*b*-PMLABe_73_-based NPs after 4h of incubation, only 40% of HepaRG cells were positive for these pegylated NPs. After 24h of incubation, all HepaRG cells had internalized PEG_42_-*b*-PMLABe_73_-based NPs but their mean fluorescence was much weaker that for the cells incubated with PMLABe_73_-based NPs as observed for macrophages.

While the uptake of 20 and 100 nm FluoSpheres® was in the same range of fluorescence intensities in macrophages, a three-fold higher fluorescence was observed in HepaRG cells incubated with 100 nm FluoSpheres® as compared to the mean found for those of 20 (**Figure 2D**). Interestingly, the means of fluorescence were also higher in macrophages than in HepaRG cells with both FluoSpheres®.

Together, these data demonstrated that the uptake of PMLABe_73_-, PEG_42_-*b*-PMLABe_73_-based NPs was faster in macrophages than in HepaRG cells although the overall internalization of PMLABe_73_-based NPs in hepatic cells at 24h was in the same range than in macrophages. Moreover, the uptake of 100nm FluoSpheres® in HepaRG cells was more efficient than for smaller particles of 20nm, and strengthened the conclusion than diameters of NPs >100 nm favored the internalization in HepaRG cells. Finally, high density of PEG on NPs (100% of PEG_42_-*b*-PMLABe_73_-derived NPs) strongly reduced NP’s uptake in both cell types.

### 3.2 Influence of PEG density on NP’s uptake by macrophages and HepaRG cells

We next decided to study the influence of various PEG densities on the surface of PMLABe-based NPs on their uptake by HepaRG cells and human macrophages. For that purpose, we mixed different proportions of the two polymers PMLABe_73_ and PEG_42_-*b*-PMLABe_73_ to prepare two additional batches of NPs: a batch of NPs prepared from a mixture of 75 wt% of PMLABe_73_ and 25 wt% of PEG_42_-*b*-PMLABe_73_ (PEG_42_-*b*-PMLABe_73_/PMLABe_73_

25/75%) and another batch of NPs prepared from a mixture of 50 wt% of PMLABe_73_ and 50 wt% of PEG_42_-*b*-PMLABe_73_(PEG_42_-*b*-PMLABe_73_/PMLABe_73_ 50/50%). Both macrophages (**Figure 3A**) and HepaRG cells (**Figure 3B**) were incubated for various times with the PMLABe_73_(100%)-, PEG_42_-*b*-PMLABe_73_/PMLABe_73_(25/75%)-, PEG_42_-*b*-PMLABe_73_/PMLABe_73_(50/50%)- and PEG_42_-*b*-PMLABe_73_(100%)-based NPs and cell uptake was quantified by flow cytometry.

**Figure 3:**
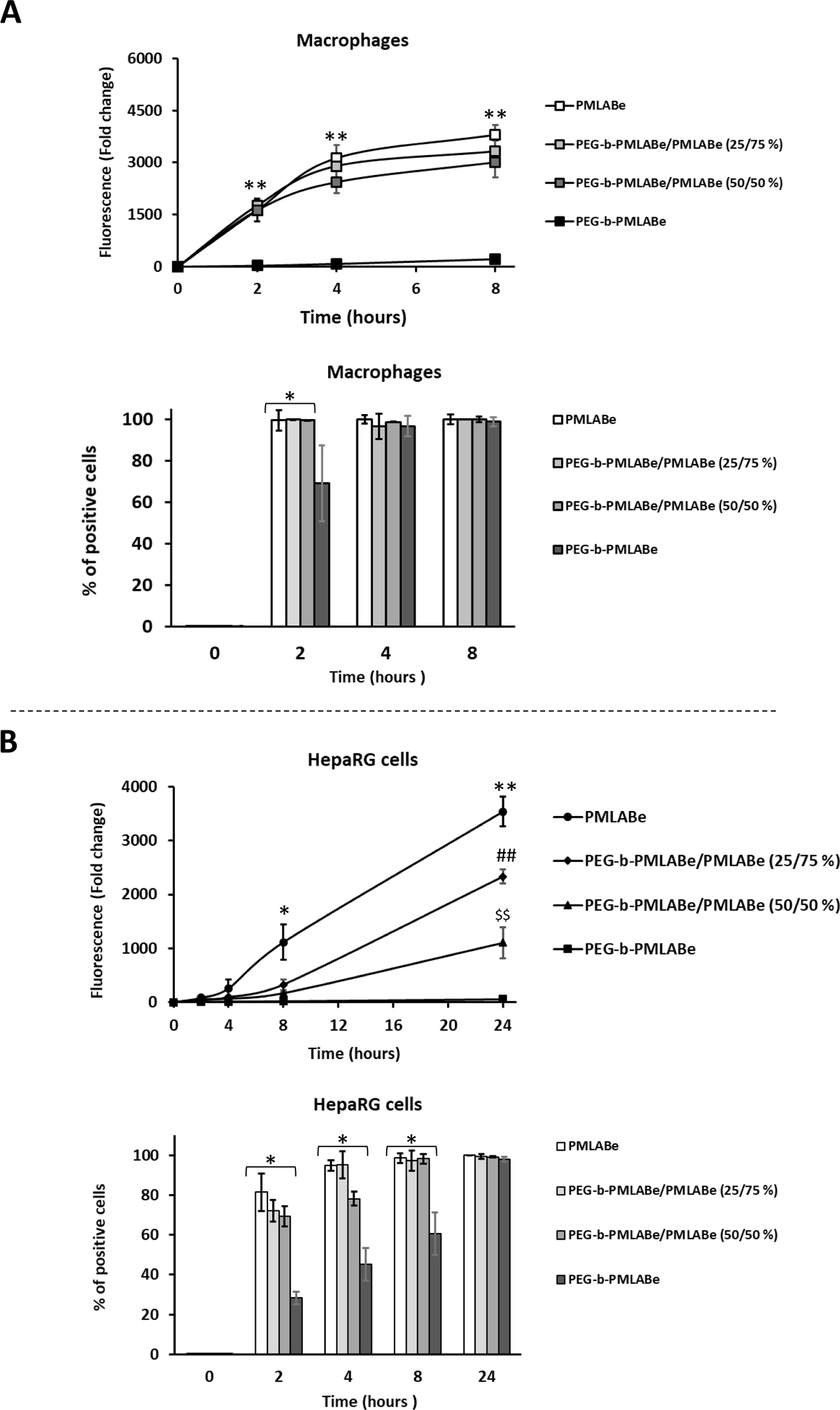
Quantification of the uptake of PMLABe_73_- and PEG_42_-b-PMLABe_73_-based NPs with various PEG densities on NP’s surface in primary macrophages and HepaRG cells by flow cytometry. Macrophages from 4 different donors (A) and HepaRG cells (B) were incubated with NPs prepared from PMLABe_73_ homopolymer and PEG_42_-*b*-PMLABe_73_ copolymer and NPs formulated with various amounts of these first two (co)polymers: PEG_42_-*b*-PMLABe_73_/PMLABe_73_ (25/75%) and a mixture of 50 wt% of PEG_42_-*b*-PMLABe_73_ and PMLABe_73_ (50/50%). Both macrophages (A) and HepaRG cells (B) were incubated for various times with these NPs and cell uptake was quantified by flow cytometry (FACSCalibur analyzer, Becton Dickinson). The curves represent the mean of fluorescence expressed in fold change of the background fluorescence measured in non-incubated cells. The histograms represent the percentage of positive macrophages (A) and HepaRG cells (B). Three independent cultures of HepaRG were performed to measure the uptake in 6 to 9 independent culture wells. * *p* < 0.05, **/##/$$ *p* < 0.01 between data obtained with cells incubated with NPs formulated with PMLABe_73_., PEG_42_-*b*-PMLABe_73_/PMLABe_73_ (25/75%) and (50/50%) versus cells incubated with PEG_42_-*b*-PMLABe_73_(100%)-based NPs.

Our data showed that macrophages incubated with PMLABe_73_(100%)-, PEG_42_-*b*-PMLABe_73_/PMLABe_73_(25/75%)-, PEG_42_-*b*-PMLABe_73_/PMLABe_73_(50/50%)-based NPs, presented similar percentages of positive cells and identical means of fluorescence (**Figure 3A)**. In contrast, incubation of macrophages with PEG_42_-*b*-PMLABe_73_(100%)-based NPs resulted in significantly lower fluorescence levels than those observed with the other three types of NPs, as previously observed in our study (**Figure 1)**. Conversely, the uptake of PMLABe_73_(100%)-, PEG_42_-*b*-PMLABe_73_/PMLABe_73_(25/75%)-, PEG_42_-*b*-PMLABe_73_/PMLABe_73_(50/50%)- and PEG_42_-*b*-PMLABe_73_(100%)-based NPs in HepaRG cells was inversely correlated to the proportion of PEG (**Figure 3B)**.

Together, these data demonstrated that only dense PEG corona on the surface of PMLABe NPs significantly inhibits the NP’s uptake by macrophages while a density of PEG as low as 25% has a strong impact on NP’s internalization in HepaRG cells.

### 3.3 In vitro cell uptake of peptide-functionalized PMLABe_73_ and PEG_42_-b-PMLABe_73_-based NPs by peripheral blood monocytic cells and macrophages

A major objective of this study was to investigate the influence of the functionalization of PMLABe-derived NPs by the peptide GBAV10-9, which was shown to exhibit a strong hepatotropism on human hepatoma cells [Vène et al., 2022]. Our goal was to determine whether the grafting of GBVA-10-9 onto PMLABe-derived NPs could affect the NP’s uptake in blood cells, hepatic HepaRG cells and macrophages. Considering the results obtained in this study regarding the effect of high PEG density on the uptake of pegylated PMLABe-derived NPs, we have chosen to engraft 10% of GBVA-10-9 peptide on PMLABe-derived NPs (**Table 1**). The engraftment was performed after NPs formulation using GBVA10-9 C-terminated with a thiol group (GBVA10-9-SH) reacting with maleimide-functionalized NPs suspensions (**Supporting information 1B**) [PMLABe_73_(90wt%)/Mal-PMLABe_73_(10wt%) or PMLABe_73_(90wt%)/Mal-PEG_62_-*b*-PMLABe_73_(10wt%)] to produce PMLABe_73_(90wt%)/GBVA10-9-PMLABe_73_(10wt%) and PMLABe_73_(90wt%)/GBVA10-9-PEG_62_-*b*-PMLABe_73_(10wt%), which showed slightly larger hydrodynamic diameters and higher polydispersity index than the non-functionalized NPs (**Table 1**).

Whole peripheral mononuclear cells (PBMC), CD14^-^, CD14^+^ and CD14^+^-derived macrophages obtained from three different donors were incubated with non-functionalized and GBVA10-9-decorated NPs and their uptake was studied by flow cytometry (**Figure 4A**). First, we observed that low percentages of PBMC and CD14^-^ internalized the four PMLABe-derived NPs, from 5 to 40% with large differences between donors. Interestingly, the uptake occurred mostly during the first 4h of incubation and did not significantly vary up to 72h of incubation. In addition, the fluorescence means remained very low demonstrating that these heterogenous cell populations internalized limited amounts of PMLABe-derived NPs. However, we also observed that GBVA10-9-PMLABe NPs were significantly more internalized in two out of the three donors (**Figure 4A**).

**Figure 4:**
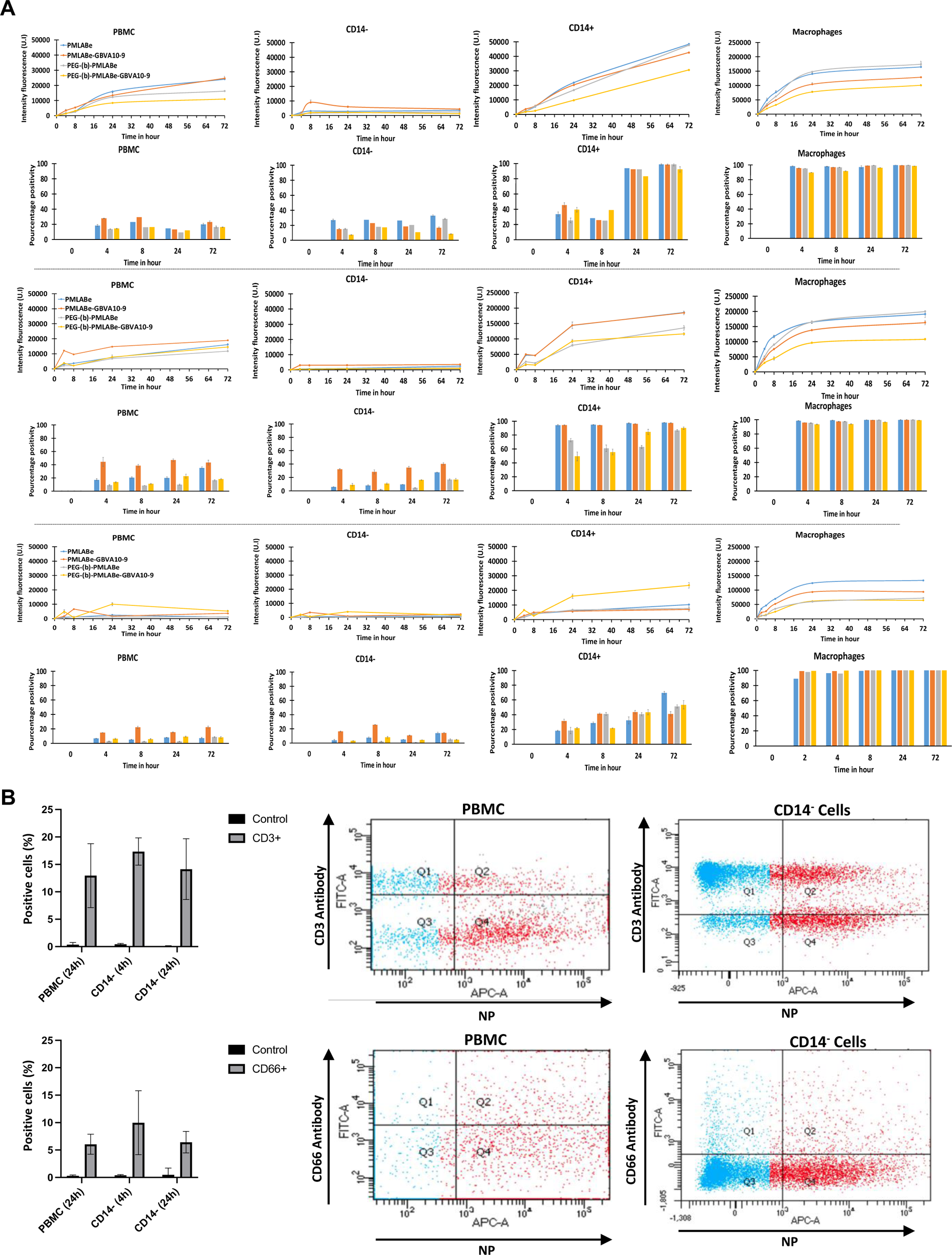
Quantification of the uptake of PMLABe_73_-, PEG_42_-b-PMLABe_73_-based NPs and their GBVA10-9 functionalized derived NPs by flow cytometry in whole PBMC populations, CD14^-^, CD14^+^ cells and primary macrophages. These experiments were performed using a LSRFortessa™ X-20 cytometer (Becton Dickinson) and data were analyzed using FACSDiva^TM^ software (Becton Dikinson). A) The curves and histograms represent the mean of fluorescence and the percentage of positive cells, respectively, for each cell types (PBMC, CD14^-^, CD14^+^ cells and macrophages) measured at 4, 8, 24 and 72h expressed in arbitrary units (A.U.) after setting up the background fluorescence at 60 (A.U.) for all cell types in each experiment. B). Percentages of DiD-Oil positive CD3^+^ and CD66^+^ cells representing the neutrophils and T lymphocytes, which had internalized DiD-Oil loaded NPs. Right: Dot plots represent typical experiments from which the chart of DiD-Oil positive CD3^+^ and CD66^+^ cells (left) were extrapolated. The data were obtained from three independent cultures of PBMC, CD14^-^, CD14^+^ cells and macrophages prepared from 3 different healthy blood donors.

Then, we quantified the uptake of the same NPs in CD14^+^ monocytes and CD14^+^-derived macrophages and found that the uptake of the four NPs was much higher in monocytes compared to PBMC with important quantitative differences between donors in terms of percentages of positive cells (from 40 to 100%) and means of fluorescence confirming large interindividual uptake capacities among healthy donors. In monocytes, no differences were observed between non-functionalized and GBVA10-9-decorated NPs. This series of experiments also further demonstrated that NP’s uptake was very efficient in macrophages since nearly all macrophages were positive at 2h of incubation and with 3 to 5-fold higher means of fluorescence compared to monocytes.

In order to determine whether other populations than monocytes were able to internalize PMLABe-derived NPs in the whole PBMC populations, we labeled the lymphocytes T with anti-CD3 and neutrophils with anti-CD66 antibodies and acquired dot plots displaying PMLABe-NP’s positive cells (DiD-Oil detected with APC channel) versus CD3^+^ or CD66^+^ cells (antibodies detected with FITC channel) in flow cytometry (**Supporting information2**, **Figure 4B**). Using this double staining, we found that ∼15% and 5 to 10% of CD3^+^ and CD66^+^ positive cells had internalized NPs, respectively, without any significant differences between the four PMLABe-derived NPs. As expected, CD3^+^ and CD66^+^ positive cell populations exhibited lower DiD-Oil staining compared to monocytes and macrophages indicating a far less efficient NP’s uptake.

### 3.4 Opsonisation of PMLABe-based NPs and influence on cell uptake

It has been previously reported that the opsonization of polystyrene-FluoSpheres^®^ [Furumoto et al., 2004 ; Vène et al., 2016], silica-[Lesniak et al., 2012] and poly(β-hydroxybutyrate)- and poly(trimethylene carbonate)-b-poly(malic acid)-[Vène et al., 2016] derived NPs strongly affected the cell uptake. Given that the opsonization by human plasma proteins considerably varied between NPs, we next investigated the influence of the opsonization of non-functionalized- and GBVA10-9-PMLABe-based NPs on their uptake by human macrophages and HepaRG hepatoma cells after 24h of incubation (**Figure 5**). To address this issue, “native” or opsonized DiDoil-loaded PMLABe-based NPs or FluoSpheres^®^ were used and the cell uptake was performed by culturing the macrophages and HepaRG cells in culture medium lacking fetal calf serum (FCS). The opsonized NPs were obtained by pre-incubating the “native NPs” with human serum prior to the dilution in the culture medium for the cell uptake.

**Figure 5:**
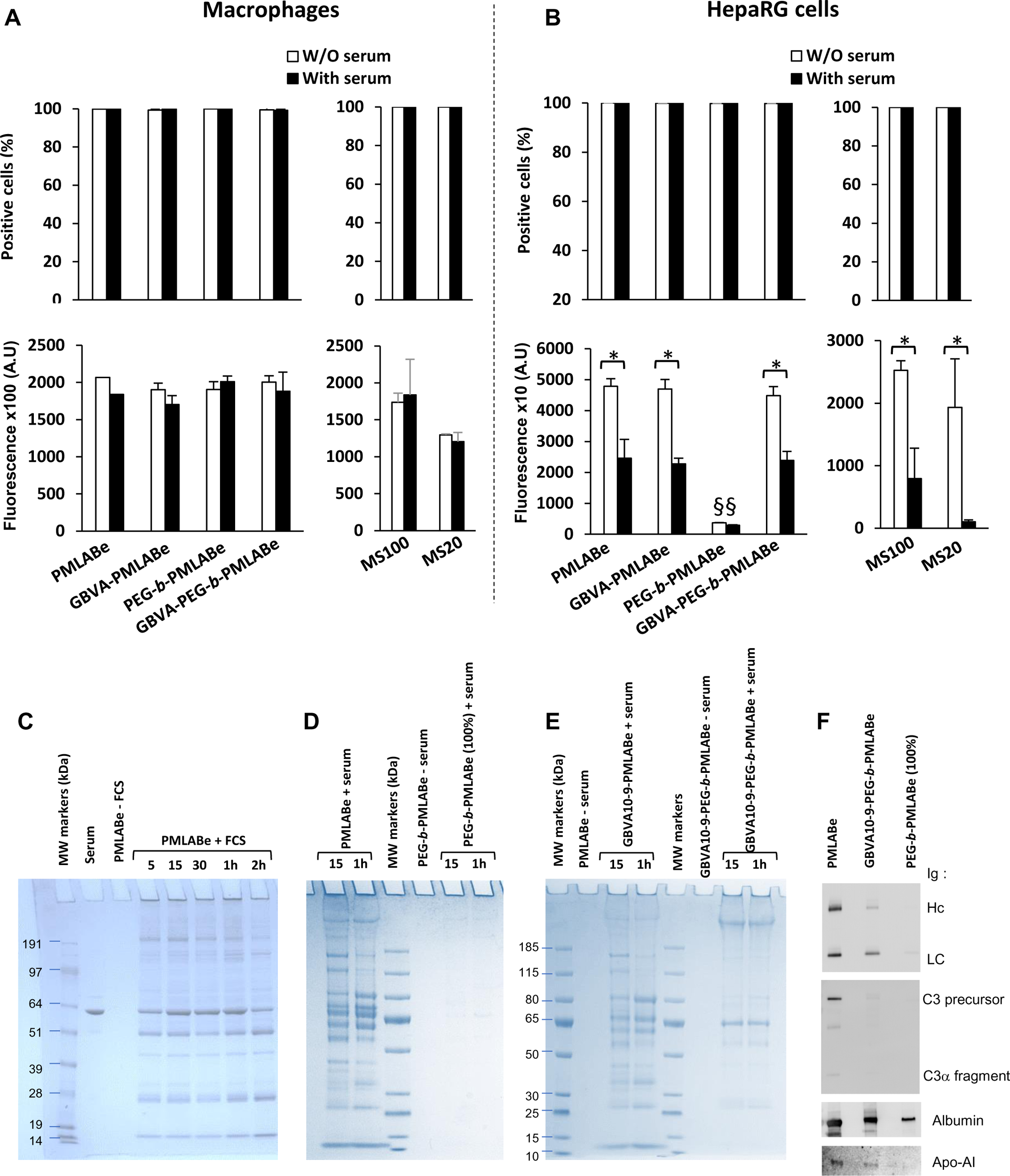
Opsonization of NPs derived from PMLABe_73_-, PEG_42_-b-PMLABe_73_- and GBVA10-functionalized-PMLABe-derived NPs by plasma proteins from human serum: Impact of opsonization on cell uptake in macrophages and HepaRG cells. Impact of serum and opsonization of the uptake of PMLABe_73_-, GBVA10-9-PMLABe_73_-, PEG_42_-b-PMLABe_73_- and GBVA10-9-PEG_42_-b-PMLABe_73_-based NPs in human macrophages (A) and HepaRG cells (B). The histograms represent the mean of fluorescence and the percentage of positive cells for measured at 24h and expressed in arbitrary units (A.U.). C) PMLABe_73_-based NPs were used to define the time-course of the opsonization assay. Incubation of NPs were performed at 5, 15, 30 min, 1 and 2h. Adsorbed proteins were loaded on SDS-PAGE and the gels were stained with Coomassie blue. Molecular weight markers (M) indicate the apparent mobility range after electrophoresis. Serum (input 1:1000 dilution, 5 μL loaded), control NPs were not incubated with serum (-serum). D) and E) Qualitative analysis for serum proteins adsorbed on PMLABe_73_-, PEG_42_-b-PMLABe_73_- and GBVA10-PEG_42_-b-PMLABe_73_-based NPs following incubation for 15min and 1h of incubation with human serum. F) Immunodetection by western blotting of immunoglobulin (heavy chains: HC ; light chains: LC), complement C3, albumin, and apolipoprotein-AI adsorbed on PMLABe_73_-, PEG_42_-b-PMLABe_73_- and GBVA10-PEG_42_-b-PMLABe_73_-based NPs.

The cell uptake by the macrophages was very efficient for all the non-functionalized- and GBVA10-9-PMLABe-based NPs with at least 95% of positive cells (**Figure 5A**). Interestingly, the mean of fluorescence in macrophages was not significantly affected by the pre-incubation of the NPs or FluoSpheres^®^ microspheres with human serum when compared to the cell uptake of “native” NPs. In addition, the deprivation in FCS did not affect this cell uptake since the overall values of fluorescence in these experiments were very similar to those found in presence of 10% FCS (**Figure 5A**).

As observed in macrophages, the uptake of non-functionalized- and GBVA10-9-PMLABe-based NPs and FluoSpheres^®^ was also very efficient in HepaRG cells with more than 95% of positive cells (**Figure 5B**). However, the opsonization of non-functionalized- and GBVA10-9-PMLABe-based NPs significantly reduced the mean of fluorescence with a 2-fold decrease for PMLABe_73-,_ PMLABe_73_/GBVA10-9-PMLABe_73_- and PMLABe_73_/GBVA10-9-PEG_62_-*b*-PMLABe_73_-NPs while internalization of PEG_62_-*b*-PMLABe_73_-NPs remained always low. Similarly, the means of fluorescence were strongly decreased for HepaRG cells incubated with opsonized FluoSpheres^®^ beads (**Figure 5B**).

The opsonization of the PMLABe_73_,(100%)-, PMLABe_73_(90%)/GBVA10-9-PMLABe_73_(10%)- and PMLABe_73_(90%)/GBVA10-9-PEG_62_-*b*-PMLABe_73_(10%)-based NPs by human serum proteins was evaluated in a cell-free protein adsorption assay and compared to that of NPs formulated with PEG_62_-*b*-PMLABe_73_(100%) only to determine if high density of PEG reduced the binding of plasma proteins (**Figure 5C-E**). In a first experiment, the protein adsorption onto PMLABe_73_,(100%)-based NPs was evaluated over a 2h time-course (**Figure 5C**). The proteins bound to the NPs were separated by electrophoresis and visualized by coomassie blue staining of polyacrylamide gels. Multiple proteins were bound onto NPs as early as 5 min after incubation with serum, and their abundance was not significantly modified with the incubation time. These two intense bands at ∼25 and 55 kDa most likely corresponded to heavy and light chains of the immunoglobulins and the abundant plasma protein with an apparent mobility weight at ∼64 kDa could be the albumin. Interestingly, NPs prepared with 100% of PEG_62_-*b*-PMLABe_73_ block copolymer showed very low binding to plasma proteins (**Figure 5D**) confirming that PEG at high density strongly reduced opsonization.

A similar procedure was used to study the opsonization of PMLABe_73_/GBVA10-9-PMLABe_73_- and PMLABe_73_/GBVA10-9-PEG_62_-*b*-PMLABe_73_-derived NPs (**Figure 5E**). The NPs prepared with PMLABe_73_/GBVA10-9-PMLABe_73_ polymers generated an opsonization nearly identical to that observed for PMLABe_73_-based NPs (**Figure 5D**). In contrast, NPs prepared with PMLABe_73_/GBVA10-9-PEG_62_-*b*-PMLABe_73_-derived NPs showed a weakest opsonization with bands that were less intense compared to the adsorption found with the other PMLABe-based NPs (**Figure 5E**).

The evaluation of the protein adsorption by coomassie blue staining of polyacrylamide was completed by immunoblotting of plasma proteins adsorbed onto PMLABe_73_-, PMLABe_73_/GBVA10-9-PEG_62_-*b*-PMLABe_73_- and PEG_62_-*b*-PMLABe_73_(100%)-derived NPs (**Figure 5F**). Specific antibodies were used to detect the human immunoglobulins, complement C3, albumin and Apolipoprotein AI. As expected, elevated amounts in immunoglobulins, complement C3, albumin and Apolipoprotein AI was found in samples of NPs formulated with PMLABe_73_ homopolymer. In contrast, NPs prepared with PEG_62_-*b*-PMLABe_73_(100%) copolymer showed much weaker signals for immunoglobulins, complement C3 and Apolipoprotein AI although albumin was found absorbed onto these NPs. Immunoblotting experiments using PMLABe_73_/GBVA10-9-PEG_62_-*b*-PMLABe_73_-derived NPs evidenced that bands obtained for immunoglobulins and complement C3 were less intense while albumin and Apolipoprotein AI were still detectable.

Together, these data indicated that NPs derived from PMLABe homopolymer were heavily opsonized, while the binding of plasma proteins was much weaker on PEG_62_-*b*-PMLABe_73_-derived NPs. This supported the conclusion that the hydrophilicity and the steric hindrance generated by the PEG block from the amphiphilic PEG_62_-*b*-PMLABe_73_ copolymer reduced the opsonization of the obtained NPs. All these data also demonstrated that the uptake of non-functionalized- and GBVA10-9-PMLABe-based NPs is differently affected by the opsonization in macrophages and HepaRG hepatoma cells. Indeed, while the opsonization of these NPs had little effect on the uptake by macrophages, the adsorption of plasmatic proteins on PMLABe-based NPs significantly affected the NP’s accumulation in HepaRG cells.

It has been demonstrated that some NPs activated the inflammasome resulting in the production of pro-inflammatory cytokines [Baron et al., 2015]. In order to determine non-functionalized- and GBVA10-9-PMLABe-based NPs may activate the production of pro-inflammatory cytokines, macrophages and HepaRG cells were incubated overnight with NPs and pro-inflammatory cytokines were quantified by ELISA assay (**Figure 6**).

**Figure 6:**
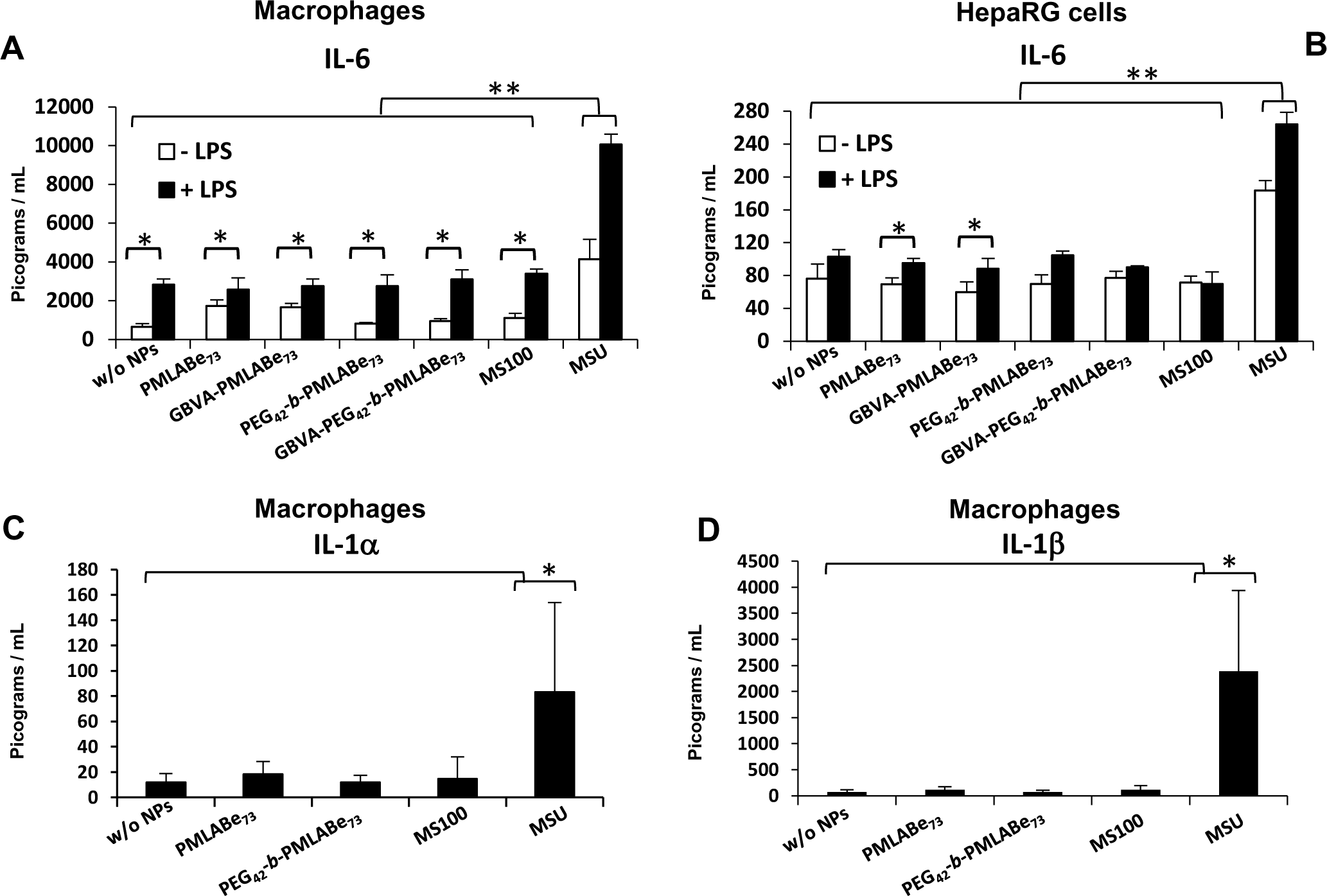
Cytokine productions in human macrophages and HepaRG cells incubated with PMLABe_73_- and PEG_42_-b-PMLABe_73_-based NPs. Interleukin-6 (IL-6), interleukin-1alpha (IL-1α) and interleukin-1bëta (IL-1β) were quantified by ELISA assay in culture media of macrophages (A, C, D) and HepaRG cells (B) following incubation with PMLABe_73_- and PEG_42_-b-PMLABe_73_-based NPs without or with functionalization by peptide GBVA10-9. Histograms of cytokine concentrations in culture media of macrophages and HepaRG cells in absence (white bars) or presence (dark bars) of the inflammasome priming factor LPS. Two different donors of monocyte-derived macrophages with n = 6 to 8 independent culture wells, 3 independent HepaRG cell culture with n = 6 to 9 independent culture wells. * *p* < 0.05, ** *p* < 0.01.

Macrophages and HepaRG cells were cultured in the absence or presence of LPS for inflammasome priming, and monosodium urate (MSU) crystals were used as positive control of sustained inflammasome activation [Gicquel et al., 2015]. The inflammation was evaluated by measuring the concentration of the pro-inflammatory cytokines IL-6 (**Figure 6A, B**) and IL-1α (**Figure 6C**) and the level of inflammasome activation was studied by determining the secretion of IL-1β (**Figure 6D**) in culture media of macrophages. In control macrophage cultures in absence of priming, IL-6 was the only detectable cytokine in culture media. The priming with LPS slightly increased the secretion of the three cytokines but the incubation with non-functionalized- and GBVA10-9-PMLABe-based NPs did not enhance the secretion of pro-inflammatory cytokines demonstrating that these NPs did not activate the inflammasome in primary macrophages. In contrast, the treatment with MSU strongly triggered the inflammasome activation visualized by the increase in cytokine secretion, as previously reported [Gicquel et al 2015]. The HepaRG cells produced much lower amounts of inflammation mediators and the IL6 was the only cytokine detectable in the culture medium. As observed for macrophages, priming with LPS slightly increased the secretion of IL-6 and the incubation with NPs did not significantly enhanced the secretion of IL-6 confirming that PMLABe-based NPs did not activate the inflammasome in this hepatocyte-like model.

### 3.5 Uptake of PMLABe-based NPs in a coculture model of hepatic cells and macrophages

Our experiments have demonstrated higher internalizations of PMLABe-based NPs in human macrophages in primary culture compared to those measured in hepatic cells using separated *in vitro* models. In order to further compare the uptake of non-functionalized- and GBVA10-9-PMLABe-based by hepatic HepaRG cells and macrophages, we set up a coculture *in vitro* model associating both HepaRG cells and human macrophages in the same culture wells, which evaluates the cell competition for the internalization of PMLABe-based NPs. To discriminate the two cell types by flow cytometry, we used HepaRG cells that stably express GFP proteins (**Supporting information 3**, **Figure 7A**). Cocultures were incubated with PMLABe_73_-, PMLABe_73_/GBVA10-9-PMLABe_73_-, PMLABe_73_/PEG_62_-*b*-PMLABe_73_ and PMLABe_73_/GBVA10-9-PEG_62_-*b*-PMLABe_73_-based NPs for 4, 8 and 24h and the red fluorescence emitted by DID-Oil loaded NPs was measured in GFP^+^ HepaRG cells and GFP^-^ negative macrophages (**Figure 7A-B**).

**Figure 7:**
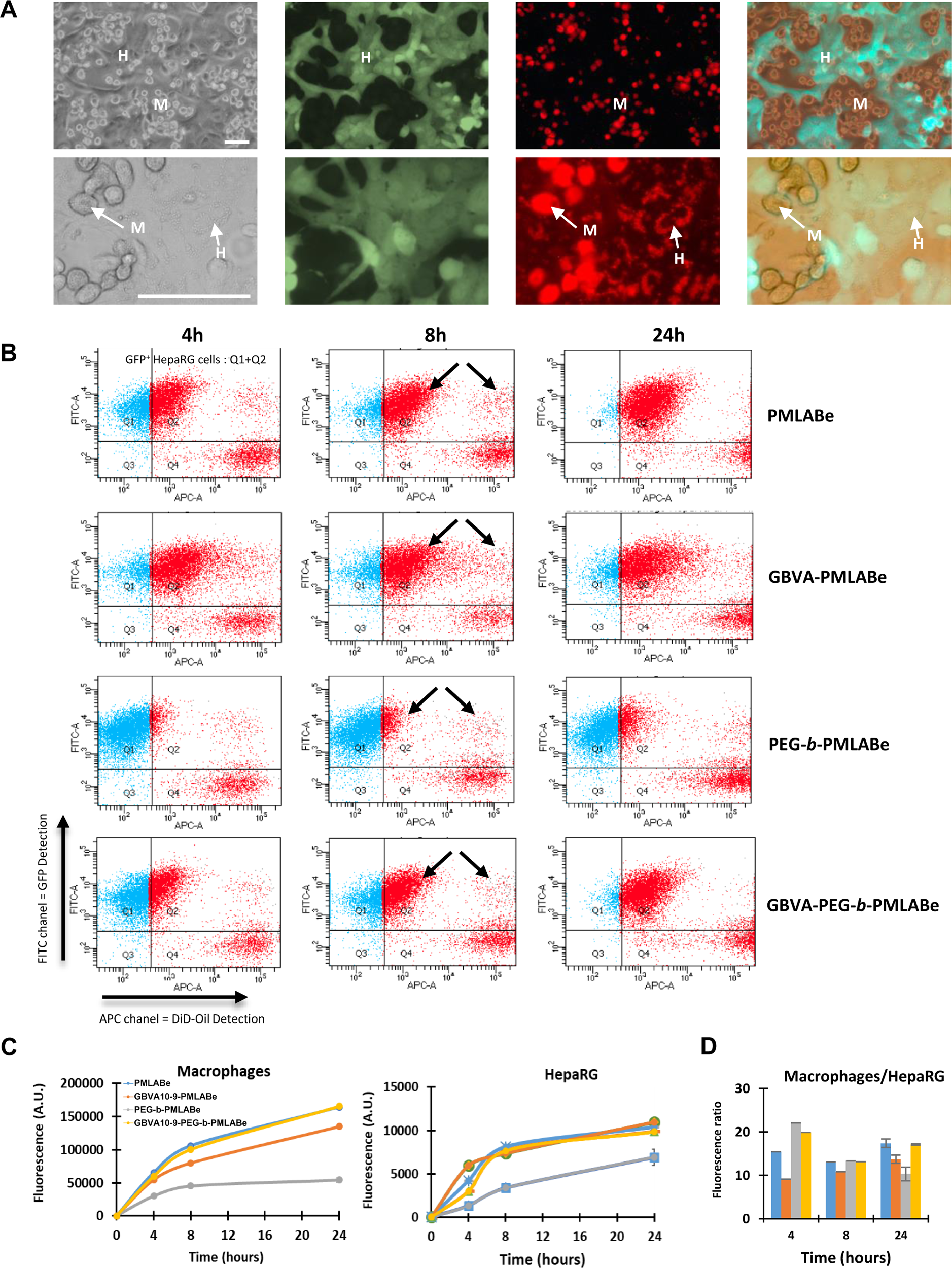
Coculture between human primary macrophages and HepaRG cells. Human macrophages (M) and GFP-expressing HepaRG cells (H) were cultured together in same well during 2 days prior to the incubation with PMLABe_73_-, GBVA10-9-PMLABe_73_-, PEG_42_-b-PMLABe_73_- and GBVA10-9-PEG_42_-b-PMLABe_73_-based NPs. Cellular uptake of fluorescent NPs loaded with DiD-Oil fluorophore was evaluated by fluorescence microscopy (A) and flow cytometry (B,C, D). A) Human macrophage (M) were visualized by phase contrast and GFP expressing HepaRG cells (H) are detected by fluorescence microscopy. Bar: 50 μm. B) Representative flow cytometry dot plots. The rectangular quadrants separate GFP-expressing HepaRG cells (Q1+Q2, FITC high) and macrophages (Q3+Q4, FITC low). The uptake of DiD-Oil loaded NPs was quantified by measuring fluorescence intensity on APC channel. C) Data are presented as mean of fluorescence intensity (A.U.) on APC channel. D) Ratio between the fluorescence intensities measured in macrophages and the fluorescence means in HepaRG cells.

The flow cytometry data showed that dot plots (FITC versus APC) allowed to discriminate GFP expressing HepaRG cells and GFP negative macrophages (**Supporting information 3**, **Figure 7B**). We confirmed that uptake of all PMLABe-derived NPs by macrophages was very efficient, these cells being far brighter than HepaRG cells at the different time points (**Figure 7A, B**). Our results also confirmed in this coculture model that the PMLABe_73_/PEG_62_-*b*-PMLABe_73_-based NPs were less internalized in both macrophages and HepaRG cells compared to the internalization of the three other NPs. Functionalization of this NP with peptide GBVA10-9 strongly increased the uptake in both cell types. Interestingly, we observed that a fraction of HepaRG cells showed an intense DiD-Oil labelling quantitatively similar to the staining in macrophages while most of HepaRG cells exhibited a ∼much weaker DiD-Oil fluorescent signal. When considering the means of fluorescence in the whole hepatic and macrophages populations, we found that the ratio of fluorescence signals emitted by DID-Oil loaded NPs in macrophages and HepaRG cells showed slight differences between the time points and that the grafting of peptide GBVA10-9 did not significantly improve the uptake of functionalized NPs in macrophages and HepaRG cells.

## IV DISCUSION

In our previous publications on the synthesis of PMLABe_73_ homopolymer and PEG_42_-*b*-PMLABe_73_ amphiphilic derivatives [Huang et al., 2012], the optimization in the routes of synthesis [Casajus et al., 2018] and first attempts in peptide functionalization of NPs prepared from these (co)polymers [Brossard et al., 2021 ; Vène et al., 2022, Brossard et al., 2022], we had not characterized in details the uptake of PMLABe_73-_ and PEG_42_-*b*-PMLABe_73_-based NPs by human hepatic cells, peripheral blood mononuclear cells and macrophages *in vitro*. In addition, we had not studied the influence of the PEG density at the surface of these NPs on the cell uptake. Yet, these *in vitro* data are important for our goals to develop biocompatible polymeric NPs capable of targeting the liver *in vivo* via systemic administration or hepatic artery injection in the treatment of hepatocellular carcinomas and for *ex vivo* perfusion of poor-quality liver grafts to improve their functions and consequently the performance of liver prior to transplantation [Del Turco et al., 2022].

The uptake of NPs prepared from PEG_42_-*b*-PMLABe_73_ only (100%) was significantly lower than that of non-PEGylated PMLABe_73_ NPs (PMLABe_73_ 100%) in both hepatic cells and macrophages at all exposure times. This effect is justified by the presence of a PEG crown on the surface of the pegylated NPs, in agreement with a large set of data obtained demonstrating the effect of the PEG crown on NP’s uptake *in vitro* [Zhang et al., 2002 ; Xie et al., 2007] and *in vivo* [Daou et al., 2009 ; Lipka et al., 2010 ; Albanese et al., 2010] in many different cell types. The PEG has exceptional physicochemical and biological properties [Gref et al., 2000 ; Davis, 2002 ; Harris 2003 ; Veronese and Pasut, 2005]. This macromolecule, soluble in water and different organic solvents, has been approved by regulatory authorities for clinical applications in humans. PEG is generally used to provide a hydrated steric barrier on the surface of NPs [Herold et al., 1989]. It is widely described in the literature that PEG provides to NPs stealth properties [Gref et al., 2000 ; Veronese and Pasut, 2005] by reducing opsonization by serum proteins and, therefore, the phagocytosis by monocytes, macrophages and non-parenchymal cells of the liver, thereby helping NPs to escape the reticuloendothelial system after intravenous administration [Gref et al., 2000 ; Veronese and Pasut, 2005].

To further study the effect of PEG on the uptake of PMLABe-derived NPs in HepaRG cells and macrophages, we prepared two additional batches of NPs with various PEG densities: PEG_42_-*b*-PMLABe_73_/PMLABe_73_(25/75%)- and PEG_42_-*b*-PMLABe_73_/PMLABe_73_(50/50%)-based NPs. The incubation of these different NPs with HepaRG cells demonstrated that the uptake of NPs was inversely correlated to the density of PEG present on their surface. These results are consistent with previous reports such as the effect of PEG density of polymer-coated NPs on their cell uptake [Du et al., 2041 ; Pelaz et al., 2015]. Conversely, the incubation of human macrophages with 0%, 25% and 50% pegylated PMLABe-based NPs had no significant effect on this NP’s internalization, while the incubation with PEG_42_-*b*-PMLABe_73_(100%)-NPs led at a much lower uptake compared to that observed with NPs exhibiting lower PEG densities. These observations indicated the absence of the dose-effect relationship between the percentage of PEG on the surface of the NPs and their uptake levels by human macrophages. Herein, we also provide new insights into the distribution of PMLABe_-_based NPs in blood cells by showing that PMLABe_73_-, PMLABe_73_/GBVA10-9-PMLABe_73_-, PMLABe_73_/PEG_62_-*b*-PMLABe_73_ and PMLABe_73_/GBVA10-9-PEG_62_-*b*-PMLABe_73_-based NPs did not strongly accumulated in PBMC with very low internalization in T lymphocytes and neutrophils while monocytes showed slightly higher uptake of these NPs. In addition, we further demonstrated the biocompatibility of PMLABe-derived NPs by showing that they did not trigger inflammasome activation and secretion of pro-inflammatory cytokines even in human macrophages, which efficiently internalizes these NPs.

The opsonization of NPs by serum proteins plays a key role in the uptake of NPs by the macrophages of the MPS system [Moghimi et al., 2012 ; Mortimer et al. 2014 ; Ren et al., 2019]. We verified whether the opsonization contributed to the uptake of PMLABe_73_(100%)- and PEG_42_-*b*-PMLABe_73_(100%)-based NPs. The opsonization led to a significant decrease in the uptake of PMLABe_73-_based NPs and in a much lesser extend for NPs prepared from PEG_42_-*b*-PMLABe_73_, in this cell line. Thus, opsonization of PMLABe-derived NPs by serum proteins reduced their uptake by HepaRG cells in agreement with previous studies studying other NPs in different *in vitro* models [Daou et al., 2009 ; Lipka et al., 2010 ; Albanese et al., 2010]. Similarly, the opsonization of polystyrene FluoSpheres^®^ also decreased the cell uptake. In contrast, the uptake levels of these NPs by human macrophages in the absence of serum remained unchanged compared to the internalization in presence of serum.

In most experiments of cell uptake performed *in vitro*, the NPs are incubated with cells maintained in their standard culture medium containing 5 to 10 % of FCS [Zauner et al., 2001; Lunov et al., 2011 ; Baron et al., 2015]. It is established that in these conditions, the NPs are rapidly opsonized by proteins of the FCS and that the opsonization affects the surface charges, the cell interactions and the cell uptake [Fleischer and Payne, 2015]. Previous reports have also studied the influence of the opsonization of polystyrene nanospheres on the hepatic biodistribution in perfused rat liver [Furumoto et al., 2004; Ogawara et al., 2004; Nagayama et al., 2007]. The authors also demonstrated that opsonization of polystyrene NPs by serum proteins reduced the hepatic disposition of these NPs, while a pre-coating of the nanospheres with purified human albumin prevented the binding of other plasma proteins and increased the blood circulation time. However, in this model of liver perfusion the authors did not discriminate the influence of the NPs opsonization on the cell uptake by macrophages versus hepatocytes and sinusoidal cells. In that respect, our data demonstrated that the opsonization of PMLABe_73_-based NPs also decreased the uptake by the hepatocyte-like HepaRG cells while it did not affect that of macrophages.

Together, these results strengthen the concept that development of novel copolymers and optimized NPs including grafting of targeting ligands associated with different routes of administration should minimize the clearance by MPS and improve either tumor accumulation [Yang et al., 2016] or specific homing in given organs for treatment of diseases [Herra-Barrera et al., 2023 ; Kim et al., 2024].

We recently reported that the peptide GBVA10-9 derived from George Baker (GB) Virus A showed a strong hepatotropism [Vène et al., 2022] using two cell-based assays: biotinylated peptide GBVA10-9 bound to fluorescent streptavidin or grafted onto biotin-poly(ethylene glycol)-*block*-poly(benzyl malate) (Biot-PEG-*b*-PMLABe)-derived NPs via streptavidin bridging strongly enhanced the endocytosis of both peptide-streptavidin conjugates and NPs in hepatoma cells [Vène et al., 2022]. This functionalization showed, however, some drawbacks incompatible with further development of clinical applications especially the use of a complex and expensive biotin-streptavidin bridging scaffold, which also strongly affected the physicochemical features of the NPs. The present study aimed at investigating the impact of the NP’s functionalization by the GBVA10-9 peptide on the cell uptake in both HepaRG cells and macrophages with a simple method of peptide grafting. We used a post-formulation strategy of peptide grafting based on the Michael reaction between a thiol group located at the C-terminal end of the peptide and a maleimide-modified PMLABe_73_ homopolymer and PEG_62_-*b*-PMLABe_73_ amphiphilic copolymer according to a recent procedure reported by our laboratory [Brossard et al., 2022].

This strategy allowed us to compare the uptake of both non-functionalized or GBVA10-9 peptide decorated PMLABe-derived NPs exhibiting only 10% (molar equivalent) in hepatic cells and macrophages. In this context, we used monocultures of HepaRG cells and human macrophages and a coculture system combining the two cell types to determine whether the functionalization of PMLABe_73_ and PEG_62_-*b*-PMLABe_73_-based NPs with GBVA10-9 peptide could favor the uptake by hepatic cells over that of macrophages. We hypothesized that GBVA10-9 peptide could enhance the uptake of functionalized PMLABe-derived NPs based on our report showing that GBVA10-9 peptide-streptavidin conjugates was more efficiently internalized by hepatoma cells than in human macrophages [Vène et al., 2022]. While the grafting of GBVA10-9 on PMLABe_73_-based NPs did not significantly affect the uptake by both HepaRG cells and macrophages, internalization of PMLABe_73_/GBVA10-9-PEG_62_-*b*-PMLABe_73_-based NPs was much higher than that of PMLABe_73_/PEG_62_-*b*-PMLABe_73_-based NPs in HepaRG cells and macrophages. In addition, the overall uptake of PMLABe_73_/GBVA10-9-PEG_62_-*b*-PMLABe_73_-based NPs reached the internalization levels observed with PMLABe_73_-based NPs. Together, these data demonstrated that the functionalization of PMLABe_73_ and PEG_62_-*b*-PMLABe_73_-based NPs by GBVA10-9 peptide, at least with this peptide density at the surface of the NPs, did not allow a better targeting of hepatocyte-like cells *in vitro*. Nevertheless, the present results reinforce our previous reports regarding the biocompatibility of PMLABe-derived NPs and further support their use to target efficiently both macrophages and hepatocytes. We are currently investigating the application of PMLABe-derived NPs in *ex vivo* perfusion of whole liver in the perspective of regenerative medicine for damaged liver grafts and in murine models of hepatocellular carcinomas.

## Author Contributions

Conceptualization, H.N., N.L., S.C.-M. and P.L.; methodology, E.V., C.R., and P.L.; software, E.V., H.N., P.L.; validation, N.L., S.C-M. and P.L.; formal analysis, H.N., N.L., S.C-M. and P.L.; investigation H.N., S.S., P.M., C.R., and P.L.; data curation, S.C.-M. and P.L..; writing—original draft preparation, H.N., S.S., S.C.-M. and P.L.; writing—review and editing, N.L., H.N., S.C.-M. and P.L.; supervision, P.L.; project administration, P.L.; funding acquisition, S.C-M. and P.L. All authors have read and agreed to the published version of the manuscript.

## Institutional Review Board Statement

Human primary monocytes were purified from buffy coat of healthy donors obtained from the Etablissement Français du Sang (EFS Rennes, France) in compliance with the French legislation on blood donation and blood products’ use and safety.

## REFERENCES

Albanese, A., Sykes, E. A. & Chan, W. C. W. Rough around the edges: the inflammatory response of microglial cells to spiky nanoparticles. ACS Nano, 2010 4, 2490–2493.

Aldosari, B. N., Alfagih, I. M., & Almurshedi, A. S. Lipid Nanoparticles as Delivery Systems for RNA-Based Vaccines. Pharmaceutics, 2021, 13(2), 206. 10.3390/pharmaceutics13020206.

Almeida DRS, Gil JF, Guillot AJ, Li J, Pinto RJB, Santos HA, Gonçalves G. Advances in Microfluidic-Based Core@Shell Nanoparticles Fabrication for Cancer Applications. Adv Healthc Mater. 2024, e2400946. doi: 10.1002/adhm.202400946.

Alving CR, Steck EA, Chapman WL Jr, Waits VB, Hendricks LD, Swartz GM Jr, Hanson WL. Therapy of leishmaniasis: superior efficacies of liposome-encapsulated drugs. Proc Natl Acad Sci U S A. 1978 Jun;75(6):2959–63. doi: 10.1073/pnas.75.6.2959.

Anselmo, A. C., & Mitragotri, S. Nanoparticles in the clinic: An update post COVID-19 vaccines. Bioengineering & Translational Medicine, 2021, 6(3), e10246. 10.1002/btm2.10246

Arvizo, R.R., Miranda, O.R., Moyano, D.F., Walden, C.A., Giri, K., Bhattacharya, R., Robertson, J.D., Rotello, V.M., Reid, J.M., Mukherjee, P. Modulating pharmacokinetics, tumor uptake and biodistribution by engineered nanoparticles. PLoS One, 2011, 6, e24374.

Baron, L., Gombault, A., Fanny, M., Villeret, B., Savigny, F., Guillou, N., Panek, C., Le Bert, M., Lagente, V., Rassendren, F., Riteau, N., Couillin, I. The NLRP3 inflammasome is activated by nanoparticles through ATP, ADP and adenosine. Cell Death Dis, 2015, 6, e1629.

Bertrand N., Leroux, J.-C. The journey of a drug-carrier in the body: An anatomo-physiological perspective. J Control Rel, 2012, 161, 152-163. 10.1016/j.jconrel.2011.09.098

Bertrand, N., Grenier, P., Mahmoudi, M., Lima, E. M., Appel, E. A., Dormont, F., Lim, J.-M., Karnik, R., Langer, R., & Farokhzad, O. C. Mechanistic understanding of in vivo protein corona formation on polymeric nanoparticles and impact on pharmacokinetics. Nature Comm., 2017, 8(1), 777. 10.1038/s41467-017-00600-w

Blanco, E., Shen, H., Ferrari M., 2015. Principles of nanoparticles design for overcoming biological barriers to drug delivery. Nat. Biotech. 33, 941–951.

Brossard C, Vlach M, Vène E, Ribault C, Dorcet V, Noiret N, Loyer P, Lepareur N, Cammas-Marion S. Synthesis of Poly(Malic Acid) Derivatives End-Functionalized with Peptides and Preparation of Biocompatible Nanoparticles to Target Hepatoma Cells. Nanomaterials (Basel). 2021 Apr 9;11(4):958. doi: 10.3390/nano11040958.

Brossard C, Vlach M, Jacquet L, Vène E, Dorcet V, Loyer P, Cammas-Marion S, Lepareur N. Hepatotropic Peptides Grafted onto Maleimide-Decorated Nanoparticles: Preparation, Characterization and *In Vitro* Uptake by Human HepaRG Hepatoma Cells. Polymers (Basel). 2022 Jun 16;14(12):2447. doi: 10.3390/polym14122447.

Cammas S, Béar MM, Harada A, Guérin P, Kataoka K. New macromolecular micelles based on degradable amphiphilic block copolymers. Macromol Chem Phys 2000; 201: 355–64.

Casajus, H., Saba, S., Vlach, M., Vène, E., Ribault, C., Tranchimand, S., Nugier-Chauvin, C., Dubreucq, E., Loyer, P., Cammas-Marion, S., & Lepareur, N. Cell Uptake and Biocompatibility of Nanoparticles Prepared from Poly(benzyl malate) (Co)polymers Obtained through Chemical and Enzymatic Polymerization in Human HepaRG Cells and Primary Macrophages. Polymers, 2018, 10(11), 1244. 10.3390/polym10111244

Coelho, T., Adams, D., Silva, A., Lozeron, P., Hawkins, P. N., Mant, T., Perez, J., Chiesa, J., Warrington, S., Tranter, E., Munisamy, M., Falzone, R., Harrop, J., Cehelsky, J., Bettencourt, B. R., Geissler, M., Butler, J. S., Sehgal, A., Meyers, R. E., … Suhr, O. B. Safety and efficacy of RNAi therapy for transthyretin amyloidosis. The New England Journal of Medicine, 2013, 369(9), 819-829. 10.1056/NEJMoa1208760

Corlu, A. and Loyer, P. Culture Conditions Promoting Hepatocyte Proliferation and Cell Cycle Synchronization. Methods Mol. Biol., 2015, 1250, 27–51.

Coty JB, Eleamen Oliveira E, Vauthier C. Tuning complement activation and pathway through controlled molecular architecture of dextran chains in nanoparticle corona. Int J Pharm., 2017, 532(2):769–778. doi: 10.1016/j.ijpharm.2017.04.048.

Couvreur P, Vauthier C. Nanotechnology: intelligent design to treat complex disease. Pharm Res. 2006, 23(7):1417–50. doi: 10.1007/s11095-006-0284-8

Davis FF. The origin of pegnology. Adv Drug Deliv Rev 2002; 54: 457–8.

Dai Q, Wilhelm S, Ding D, Syed AM, Sindhwani S, Zhang Y, Chen YY, MacMillan P, Chan WCW. Quantifying the Ligand-Coated Nanoparticle Delivery to Cancer Cells in Solid Tumors. ACS Nano., 2018, 12(8):8423–8435. doi: 10.1021/acsnano.8b03900.

Daou, T. J., Li, L., Reiss, P., Josserand, V., Texier, I. Effect of Poly(ethylene glycol) Length on the in Vivo Behavior of Coated Quantum Dots. Langmuir, 2009, 25, 3040–3044.

Dawidczyk, C.M., Kim, C., Park, J.H., Russell, L.M., Lee, K.H., Pomper, M.G., Searson, P.C. State-of-the-art in design rules for drug delivery platforms: lessons learned from FDA-approved nanomedicines. J. Control. Rel., 2014, 187, 133–144.

D’Addio, S. M., Saad, W., Ansell, S. M., Squiers, J. J., Adamson, D. H., Herrera-Alonso, M., Wohl, A. R., Hoye, T. R., Macosko, C. W., Mayer, L. D., Vauthier, C., & Prud’homme, R. K. Effects of block copolymer properties on nanocarrier protection from in vivo clearance. J. Control. Rel., 2012, 162(1), 208-217. 10.1016/j.jconrel.2012.06.020

Del Turco S, Cappello V, Tapeinos C, Moscardini A, Sabatino L, Battaglini M, Melandro F, Torri F, Martinelli C, Babboni S, Silvestrini B, Morganti R, Gemmi M, De Simone P, Martins PN, Crocetti L, Peris A, Campani D, Basta G, Ciofani G, Ghinolfi D. Cerium oxide nanoparticles administration during machine perfusion of discarded human livers: A pilot study. Liver Transplant Off Publ Am Assoc Study Liver Dis Int Liver Transplant Soc., 2022, 28:1173–1185. doi: 10.1002/lt.26421.

Dolgin, E. The tangled history of mRNA vaccines. Nature, 2021, 597(7876), 318-324. 10.1038/d41586-021-02483-w

Du, X.J., Wang, J.L., Liu, W.W., Yang, J.X., Sun, C.Y., Li, H.J., Shen, S., Luo, Y.L., Ye, X.D., Zhu, Y.H., Yang, X.Z., Wang, J., 2015. Regulating the surface poly(ethylene glycol) density of polymeric nanoparticles and evaluating its role in drug delivery in vivo. Biomaterials 69, 1–11. Editorial., 2014. Time to deliver. Nat. Biotechnol. 32, 961.

Dutkowski P, Polak WG, Muiesan P, Schlegel A, Verhoeven CJ, Scalera I, DeOliveira ML, Kron P, Clavien PA. First Comparison of Hypothermic Oxygenated PErfusion Versus Static Cold Storage of Human Donation After Cardiac Death Liver Transplants: An International-matched Case Analysis. Ann Surg., 2015, 262(5):764–70; discussion 770-1. doi: 10.1097/SLA.0000000000001473.

Fattal E, Rojas J, Youssef M, Couvreur P, Andremont A. Liposome-entrapped ampicillin in the treatment of experimental murine listeriosis and salmonellosis. Antimicrob Agents Chemother. 1991, 4, 770–2. doi: 10.1128/AAC.35.4.770.

Fleischer, C.C., Payne, C.K., 2014. Nanoparticle-cell interactions: molecular structure of the protein corona and cellular outcomes. Acc Chem Res. 47, 2651–2659.

Frank, M.M., Fries, L.F. The role of complement in inflammation and phagocytosis. Immunol. Today, 1991, 12, 322–326.

Furumoto, K., Nagayama, S., Ogawara, K., Takakura, Y., Hashida, M., Higaki, K., Kimura, T., 2004. Hepatic uptake of negatively charged particles in rats: possible involvement of serum proteins in recognition by scavenger receptor. J. Control. Rel., 2004, 97, 133–141.

Gao H, Zhang Q, Yang Y, Jiang X, He Q. Tumor homing cell penetrating peptide decorated nanoparticles used for enhancing tumor targeting delivery and therapy. Int J Pharm 2015; 478: 240–50.

Gicquel, T., Robert, S., Loyer, P., Victoni, T., Bodin, A., Ribault, C., Gleonnec, F., Couillin, I., Boichot, E., Lagente, V. IL-1β production is dependent on the activation of purinergic receptors and NLRP3 pathway in human macrophages. FASEB J., 2015, 29, 4162–73.

Gref R, Lück M, Quellec P, Marchand M, Dellacherie E, Harnisch S, Blunk T, Müller RH. ’Stealth’ corona-core nanoparticles surface modified by polyethylene glycol (PEG): influences of the corona (PEG chain length and surface density) and of the core composition on phagocytic uptake and plasma protein adsorption. Colloids Surf B Biointerfaces. 2000,18(3-4):301–313. doi: 10.1016/s0927-7765(99)00156-3.

Gustafson, H. H., Holt-Casper, D., Grainger, D. W., & Ghandehari, H. Nanoparticle Uptake: The Phagocyte Problem. Nano Today, 2015, 10(4), 487-510. 10.1016/j.nantod.2015.06.006

Herold, D. A., Keil, K., Bruns, D. E. Oxidation of polyethylene glycols by alcohol dehydrogenase. Biochem. Pharmacol., 1989, 38, 73–76.

Herrera-Barrera M, Ryals RC, Gautam M, Jozic A, Landry M, Korzun T, Gupta M, Acosta C, Stoddard J, Reynaga R, Tschetter W, Jacomino N, Taratula O, Sun C, Lauer AK, Neuringer M, Sahay G. Peptide-guided lipid nanoparticles deliver mRNA to the neural retina of rodents and nonhuman primates. Sci Adv. 2023 Jan 13;9(2):eadd4623. doi: 10.1126/sciadv.add4623.

Harris, J.M., Chess, R.B., 2003. Effect of pegylation on pharmaceuticals. Nat. Rev. 2, 214–221.

Hoffman AS. The origins and evolution of “controlled” drug delivery systems. J Control Rel., 2008; 132: 153–63.

Huang, Z.W., Laurent, V., Chetouani, G., Ljubimova, J.Y., Holler, E., Benvegnu, T., Loyer, P., Cammas-Marion, S. New functional degradable and bio-compatible nanoparticles based on poly(malic acid) derivatives for site-specific anti-cancer drug delivery. Int. J. Pharm., 2012, 423, 84–92.

Ishida, T., Ichihara, M., Wang, X., Kiwada, H. Spleen plays an important role in the induction of accelerated blood clearance of PEGylated liposomes. J. Control. Rel., 2006, 115, 243–250.

Jacobs, F., Wisse, E., De Geest, B. The role of liver sinusoidal cells in hepatocyte-directed gene transfer. Am. J. Pathol., 2010, 176, 14–21.

Kim J, Eygeris Y, Ryals RC, Jozić A, Sahay G. Strategies for non-viral vectors targeting organs beyond the liver. Nat Nanotechnol. 2024 Apr;19(4):428–447. doi: 10.1038/s41565-023-01563-4.

Kola, I., Landis, J., 2004. Can the pharmaceutical industry reduce attrition rates? Nat. Rev. 3, 711–715.

Kouser L, Paudyal B, Kaur A, Stenbeck G, Jones LA, Abozaid SM, Stover CM, Flahaut E, Sim RB, Kishore U. Human Properdin Opsonizes Nanoparticles and Triggers a Potent Pro-inflammatory Response by Macrophages without Involving Complement Activation. Front Immunol., 2018, 9:131. doi: 10.3389/fimmu.2018.00131.

Kristen, A. V., Ajroud-Driss, S., Conceição, I., Gorevic, P., Kyriakides, T., & Obici, L. Patisiran, an RNAi therapeutic for the treatment of hereditary transthyretin-mediated amyloidosis. Neurodegenerative Disease Management, 2019, 9(1), 5-23. 10.2217/nmt-2018-0033

Lesniak, A., Fenaroli, F., Monopoli, M.P., Å, Dawson, K.A., Salvati, A., 2012. Effects of the presence or absence of a protein corona on silica nanoparticle uptake and impact on cell. ACS Nano 6, 5845–5857.

Lipka J, Semmler-Behnke M, Sperling RA, Wenk A, Takenaka S, Schleh C, Kissel T, Parak WJ, Kreyling WG. Biodistribution of PEG-modified gold nanoparticles following intratracheal instillation and intravenous injection. Biomaterials, 2010, (25):6574–81. doi: 10.1016/j.biomaterials.2010.05.009.

Loyer, P., Cammas-Marion, S., 2014. Natural and synthetic poly(malic acid)-based derivates: a family of versatile biopolymers for the design of drug nanocarriers. J. Drug Target. 22, 556–575.

Luan J, Ju D. Inflammasome: A doubled-edged sword in liver diseases. Front Immunol. 2018, 9, 2201. doi: 10.3389/fimmu.2018.02201.

Lunov, O., Syrovets, T., Loos, C., Nienhaus, G.U., Mailänder, V., Landfester, K., Rouis, M., Simmet, T., 2011. Amino-functionalized polystyrene nanoparticles activate the NLRP3 inflammasome in human macrophages. ACS Nano. 5, 9648–9657.

Maeda, H., Nakamura, H., Fang, J., 2013. The EPR effect for macromolecular drug delivery to solid tumors: Improvement of tumor uptake, lowering of systemic toxicity, and distinct tumor imaging in vivo. Adv. Drug Deliv. Rev., 65, 71–79.

Maeda H., Tsukigawa K., Fang J., 2016. A retrospective 30 years after discovery of the enhanced permeability and retention effect to solid tumors: next-generation chemotherapeutics and photodynamic therapy-problems, solutions, and prospects. Microcirculation, 23, 173–182.

Mahon E, Salvati A, Baldelli Bombelli F, Lynch I, Dawson KA. Designing the nanoparticles-biomolecule interface for targeting and therapeutic delivery. J Control Release 2012; 161: 164–74.

McNeil, S. E. Evaluation of nanomedicines : Stick to the basics. Nature Reviews Materials, 2016, 1(10), Article 10. 10.1038/natrevmats.2016.73

Means N, Elechalawar CK, Chen WR, Bhattacharya R, Mukherjee P. Revealing macropinocytosis using nanoparticles. Mol Aspects Med., 2022, 83:100993. doi: 10.1016/j.mam.2021.100993.

Moghimi, S. M., Hunter, A. C., Andresen, T. L. Factors controlling nanoparticle pharmacokinetics: an integrated analysis and perspective. Annu. Rev. Pharmacol. Toxicol. 2012, 52, 481–503.

Mortimer GM, Butcher NJ, Musumeci AW, Deng ZJ, Martin DJ, Minchin RF. Cryptic epitopes of albumin determine mononuclear phagocyte system clearance of nanomaterials. ACS Nano., 2014, 8(4):3357–66. doi: 10.1021/nn405830g.

Nagayama, S., Ogawara, K., Fukuoka, Y., Higaki, K., Kimura, T., 2007. Time-dependent changes in opsonin amount associated on nanoparticles alter their hepatic uptake characteristics. Int J Pharm. 342, 215–221.

Nemes B, Gámán G, Polak WG, Gelley F, Hara T, Ono S, Baimakhanov Z, Piros L, Eguchi S.. Extended-criteria donors in liver transplantation Part II: reviewing the impact of extended-criteria donors on the complications and outcomes of liver transplantation. Expert Rev Gastroenterol Hepatol. 2016, 10:841–859. doi: 10.1586/17474124.2016.1149062.

Noack K, Bronk SF, Kato A, Gores GJ. The greater vulnerability of bile duct cells to reoxygenation injury than to anoxia. Implications for the pathogenesis of biliary strictures after liver transplantation. Transplantation, 1993, 56:495–500. doi: 10.1097/00007890-199309000-00001.

O’Brien, M.E., Wigler N, Inbar M, Rosso R, Grischke E, Santoro A, Catane R, Kieback DG, Tomczak P, Ackland SP, Orlandi F, Mellars L, Alland L, Tendler C., 2004. Reduced cardiotoxicity and comparable efficacy in a phase III trial of pegylated liposomal doxorubicin HCl (CAELYX/Doxil) versus conventional doxorubicin for first-line treatment of metastatic breast cancer. Ann. Oncol. 15, 440–449.

Ogawara, K., Furumoto, K., Nagayama, S., Minato, K., Higaki, K., Kai, T., Kimura, T., 2004. Pre-coating with serum albumin reduces receptor-mediated hepatic disposition of polystyrene nanosphere: implications for rational design of nanoparticles. J. Control. Rel. 100, 451–455.

Owens, D.E., Peppas, N.A. Opsonization, biodistribution, and pharmacokinetics of polymeric nanoparticles. Int. J. Pharm., 2006, 307, 93–102.

Pati R, Shevtsov M, Sonawane A. Nanoparticle Vaccines Against Infectious Diseases. Front Immunol. 2018, 9:2224. doi: 10.3389/fimmu.2018.02224.

Pelaz B, del Pino P, Maffre P, Hartmann R, Gallego M, Rivera-Fernández S, de la Fuente JM, Nienhaus GU, Parak WJ. Surface Functionalization of Nanoparticles with Polyethylene Glycol: Effects on Protein Adsorption and Cellular Uptake. ACS Nano., 2015, 9(7):6996–7008. doi:10.1021/acsnano.5b01326.

Penn, C. A., Yang, K., Zong, H., Lim, J.-Y., Cole, A., Yang, D., Baker, J., Goonewardena, S. N., & Buckanovich, R. J. Therapeutic Impact of Nanoparticle Therapy Targeting Tumor-Associated Macrophages. Molecular Cancer Therapeutics, 2018, 17(1), 96-106. 10.1158/1535-7163.MCT-17-0688

Price, L. S. L., Stern, S. T., Deal, A. M., Kabanov, A. V., & Zamboni, W. C. A reanalysis of nanoparticle tumor delivery using classical pharmacokinetic metrics. Science Advances, 2020, 6(29), eaay9249. 10.1126/sciadv.aay9249

Raemdonck K, De Smedt SC. Lessons in simplicity that should shape the future of drug delivery. Nature Biotechnology 2015; 33: 1026–7.

Reddy, L. H., Couvreur, P., 2011. Nanotechnology for therapy and imaging of liver diseases. J. Hepatol. 55, 1461–1466.

Ren J, Cai R, Wang J, Daniyal M, Baimanov D, Liu Y, Yin D, Liu Y, Miao Q, Zhao Y, Chen C. Precision Nanomedicine Development Based on Specific Opsonization of Human Cancer Patient-Personalized Protein Coronas. Nano Lett., 2019, 19(7):4692–4701. doi: 10.1021/acs.nanolett.9b01774.

Sahay, G., Alakhova, D.Y., Kabanov, A. V. Endocytosis of nanomedicines. J. Control. Rel., 2010, 145, 182–195.

Schlegel A, de Rougemont O, Graf R, Clavien PA, Dutkowski P. Protective mechanisms of end-ischemic cold machine perfusion in DCD liver grafts. J Hepatol., 2013, 58(2):278–86. doi: 10.1016/j.jhep.2012.10.004.

Shi, N.-Q.; Li, Y.; Zhang, Y.; Shen, N.; Qi, L.; Wang, S.-R.; Qi, X.-R. Intelligent “Peptide-Gathering Mechanical Arm” Tames Wild “Trojan-Horse” Peptides for the Controlled Delivery of Cancer Nanotherapeutics. ACS Appl. Mater. Interfaces., 2017, 9, 41767–41781.

Smits LP, Coolen BF, Panno MD, Runge JH, Nijhof WH, Verheij J, Nieuwdorp M, Stoker J, Beuers UH, Nederveen AJ, Stroes ES. Noninvasive Differentiation between Hepatic Steatosis and Steatohepatitis with MR Imaging Enhanced with USPIOs in Patients with Nonalcoholic Fatty Liver Disease: A Proof-of-Concept Study. Radiology. 2016, 278(3):782–91. doi: 10.1148/radiol.2015150952.

Stinchcombe, T.E., 2007. Nanoparticle albumin-bound paclitaxel: a novel Cremphor-EL-free formulation of paclitaxel. Nanomedicine 4, 415–423.

Stylianopoulos T, Jain RK. Design considerations for nanotherapeutics in oncology. Nanomed: Nanotechnol Biol Med 2015; 11: 1893–907.

Sun, H.; Dong, Y.; Feijen, J.; Zhong, Z. Peptide-decorated polymeric nanomedicines for precision cancer therapy. J. Control. Rel., 2018, 290, 11–27.

Tacke F. Targeting hepatic macrophages to treat liver diseases. J Hepatol., 2017, 66:1300–12.

Tenzer, S., Docter, D., Kuharev, J., Musyanovych, A., Fetz, V., Hecht, R., Schlenk, F., Fischer, D., Kiouptsi, K., Reinhardt, C., Landfester, K., Schild, H., Maskos, M., Knauer, S.K., Stauber, R.H., 2013. Rapid formation of plasma protein corona critically affects nanoparticle pathophysiology. Nat. NanoTechnol. 8, 772–781.

Thioune, O., Fessi, H., Devissaguet, J. P., Puisieux, F. Preparation of pseudolatex by nanoprecipitation: influence of the solvent nature on intrinsic viscosity and interaction constant. Int. J. Pharm., 1997, 146, 233–238.

Topete A, Barbosa S, Taboada P. Intelligent micellar polymeric nanocarriers for therapeutics and diagnosis. J Appl Polym Sci, 2015, 132: DOI: 10.1002/APP.42650.

Torchilin V. P., 2006. Multifunctional nanocarriers. Adv Drug Deliv Rev, 58, 1532–1555.

van Rijn R, Schurink IJ, de Vries Y, van den Berg AP, Cortes Cerisuelo M, Darwish Murad S, Erdmann JI, Gilbo N, de Haas RJ, Heaton N, van Hoek B, Huurman VAL, Jochmans I, van Leeuwen OB, de Meijer VE, Monbaliu D, Polak WG, Slangen JJG, Troisi RI, Vanlander A, de Jonge J, Porte RJ; DHOPE-DCD Trial Investigators. Hypothermic Machine Perfusion in Liver Transplantation - A Randomized Trial. N Engl J Med., 2021, 384(15):1391–1401. doi: 10.1056/NEJMoa2031532.

Vène E, Barouti G, Jarnouen K, Gicquel T, Rauch C, Ribault C, Guillaume SM, Cammas-Marion S, Loyer P. Opsonisation of nanoparticles prepared from poly(β-hydroxybutyrate) and poly(trimethylene carbonate)-b-poly(malic acid) amphiphilic diblock copolymers: Impact on the in vitro cell uptake by primary human macrophages and HepaRG hepatoma cells. Int J Pharm. 2016 Nov 20;513(1-2):438–452. doi: 10.1016/j.ijpharm.2016.09.048.

Vène E, Jarnouen K, Ribault C, Vlach M, Verres Y, Bourgeois M, Lepareur N, Cammas-Marion S, Loyer P. Circumsporozoite Protein of *Plasmodium berghei*- and George Baker Virus A-Derived Peptides Trigger Efficient Cell Internalization of Bioconjugates and Functionalized Poly(ethylene glycol)-*b*-poly(benzyl malate)-Based Nanoparticles in Human Hepatoma Cells. Pharmaceutics. 2022, 14(4):804. doi: 10.3390/pharmaceutics14040804.

Veronese, F.M., Pasut, G., 2005. PEGylation, successful approach to drug delivery. Drug Discov Today. 10, 1451–1458.

Wicki, A., Witzigmann, D., Balasubramanian, V., Huwyler, J. Nanomedicine in cancer therapy: Challenges, opportunities, and clinical applications J. Control. Rel., 2015, 200, 138–157

Wilhelm, S., Tavares, A. J., Dai, Q., Ohta, S., Audet, J., Dvorak, H. F., & Chan, W. C. W. Analysis of nanoparticle delivery to tumours. Nature Reviews Materials, 2016, 1(5), Article 5. 10.1038/natrevmats.2016.14

Xie, J., Xu, C., Kohler, N., Hou, Y., & Sun, S. Controlled PEGylation of Monodisperse Fe3O4 Nanoparticles for Reduced Non-Specific Uptake by Macrophage Cells. Advanced Materials, 2007, 19(20), 3163-3166. 10.1002/adma.200701975

Yang B, Han X, Ji B, Lu R., 2016. Competition between tumor and mononuclear phagocyte system cause the low tumor distribution of nanoparticles and strategies to improve tumor accumulation. Curr. Drug Deliv. 13, publication ahead of printing. DOI: 10.2174.

Yao CG, Martins PN. Nanotechnology Applications in Transplantation Medicine. Transplantation, 2020, 104:682–693.

Youden, B., Jiang, R., Carrier, A. J., Servos, M. R., & Zhang, X. A Nanomedicine Structure-Activity Framework for Research, Development, and Regulation of Future Cancer Therapies. ACS Nano, 2022, 16(11), 17497-17551. 10.1021/acsnano.2c06337

Zauner, W, Farrow, NA, Haines, AM., 2001. In vitro uptake of polystyrene microspheres: effect of particle size, cell line and cell density. J Control. Rel. 71, 39–51.

Zhang Y, Kohler N, Zhang M. Surface modification of superparamagnetic magnetite nanoparticles and their intracellular uptake. Biomaterials, 2002, (7):1553–61. doi: 10.1016/s0142-9612(01)00267-8

Zhang, X., Ng, H.L., Lu, A., Lin, C., Zhou, L., Lin, G., Zhang, Y., Yang, Z., Zhang, H., 2016. Drug delivery system targeting advanced hepatocellular carcinoma: Current and future. Nanomedicine 12, 853–869.

Zhu, Y., Feijen, J., Zhong, Z. Dual-targeted nanomedicines for enhanced tumor treatment. Nano Today, 2018, 18, 65-85. 10.1016/j.nantod.2017.12.007.

