## Supplemental information for "Efficient internalization of poly(benzyl malate) and poly(ethylene glycol)-*b*-poly(benzyl malate) copolymer based nanoparticles by human hepatic HepaRG cells and macrophages : Impact of nanoparticle functionalization by GBVA10-9 peptide on cell uptake"

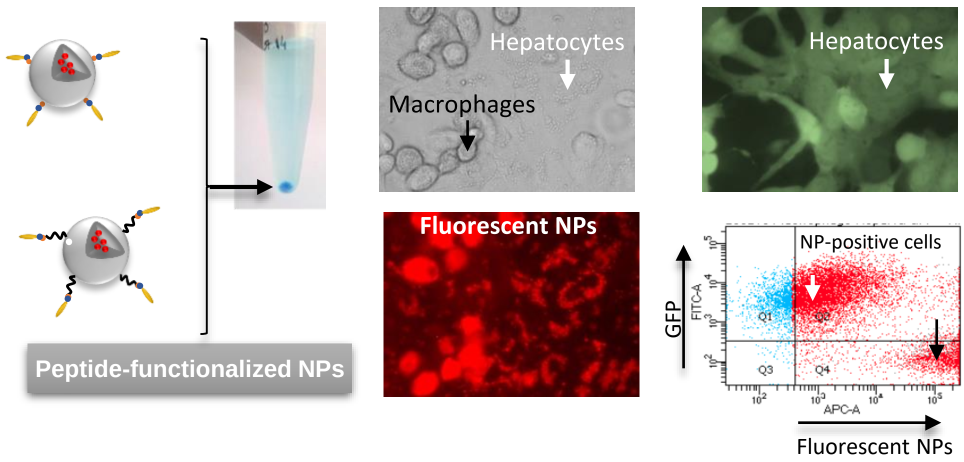


Graphical abstract

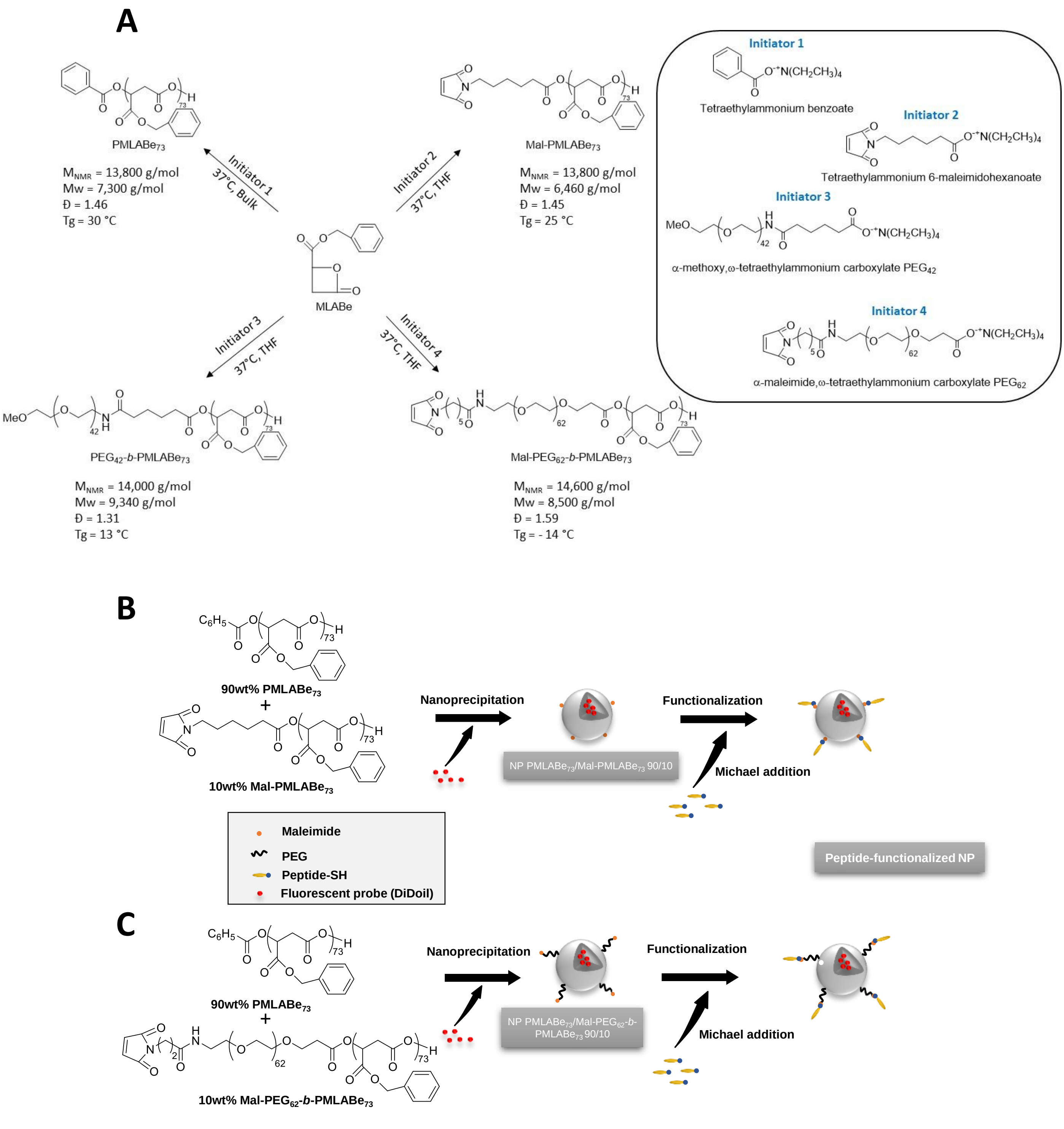


**Supporting Information 1 : Synthesis of (co)polymers, formulation and functionalizationof nanoparticles.** The synthesis of PMLABe$_{73}$, PEG$_{42}$-*b*-PMLABe$_{73}$, Mal-PMLABe$_{73}$, and Mal-PEG$_{62}$-*b*-PMLABe$_{73}$ (co)polymers was obtained through aROP of MLABe as previously reported [Brossard et al., Polymers, 2022, 14, 2247].

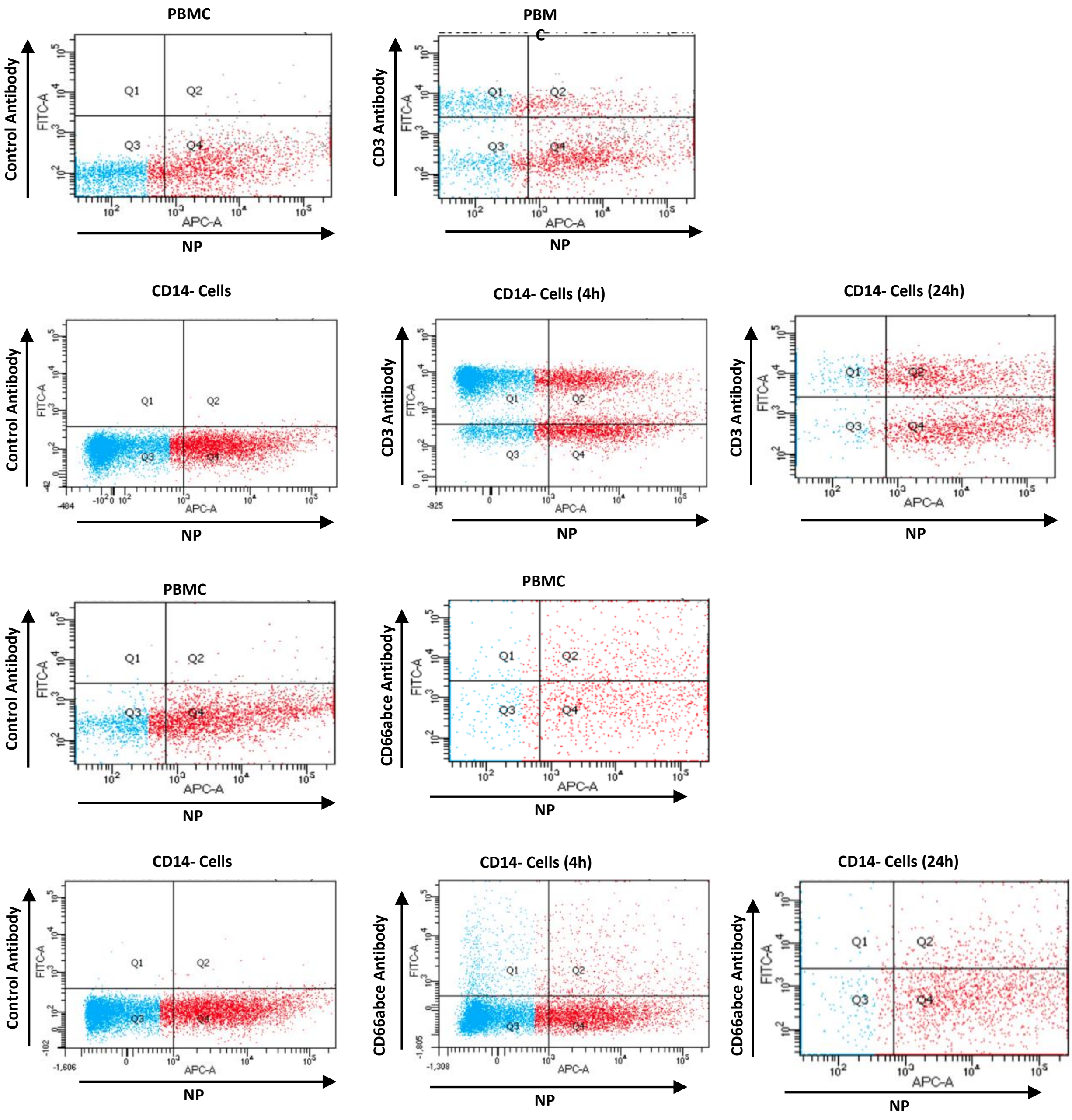


**Supporting Information 2.**

Dot plots represent typical experiments from which the chart of DiD-Oil positive CD3$^+$ and CD66$^+$ cells in figure 4 were extrapolated. Dot plots with control isotype antibodies show the gating for background fluorescence used to detect CD3$^+$ and CD66$^+$ cells using specific antibodies. Similar data were obtained in three independent cultures of PBMC, CD14$^-$, CD14$^+$ cells and macrophages prepared from 3 different healthy blood donors.

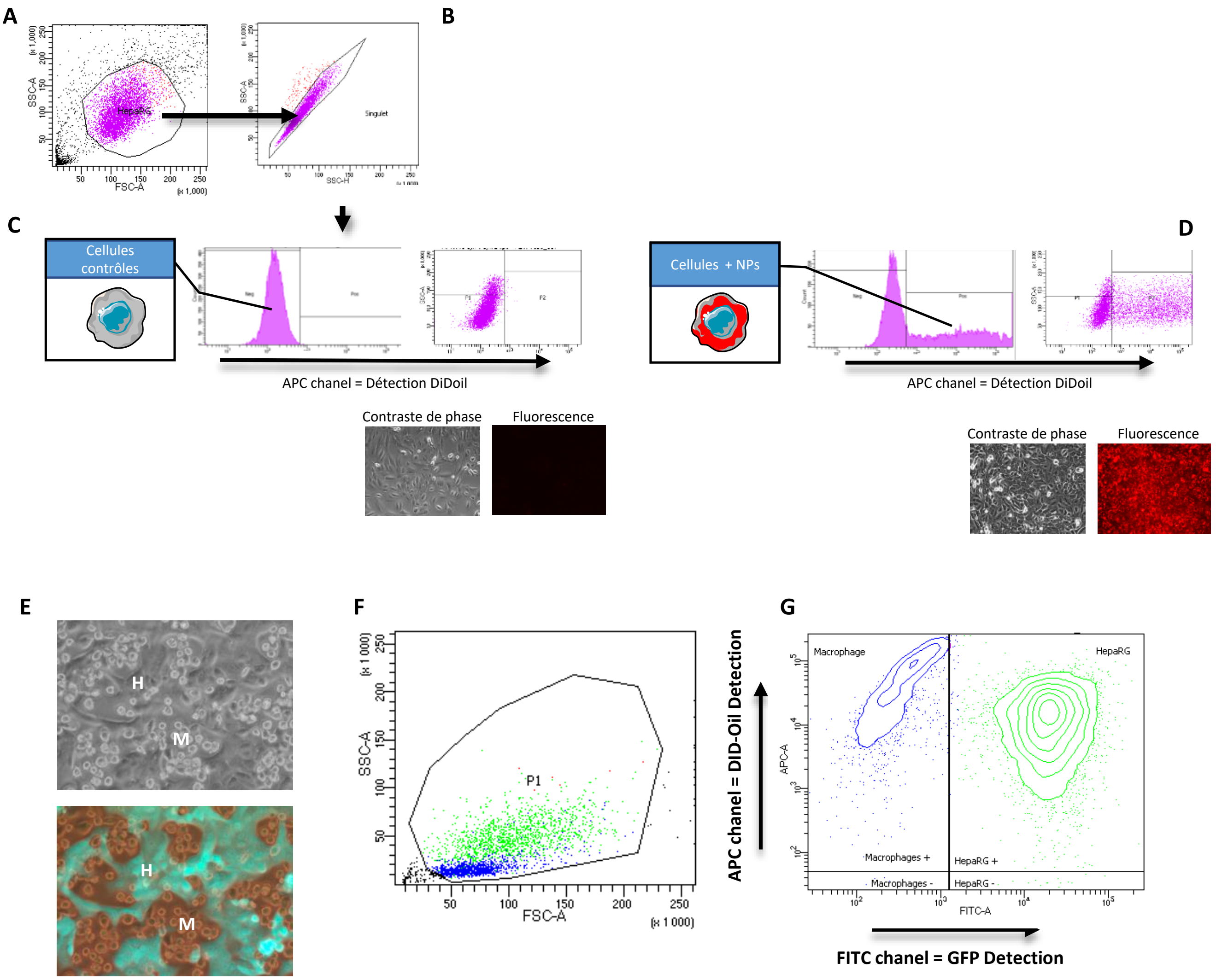


**Supporting Information 3 : Procedure cell detection and settings for flow cytometry analysis.**
The cells analyzed by flow cytometry were gated using a dot plot with the SSC (granularity: y) and FSC (size: x) parameters (**A**). Then, cells from this first population were further gated using the SSC-Height versus SSC-Area to isolate the single cells (**B**). From the single cell population, fluorescence was analyzed (**C**-upper histogram : APC fluorescence on x axis versus cell counts on y axis and/or dot plots APC fluorescence on x axis versus SSC-A on y axis) in cells that were not incubated with DiD-Oil fluorescent NPs to define the "background" fluorescence also called auto-fluorescence of negative cells (Neg/P1). Then, the cells incubated with the different fluorescent NPs were analyzed (**D**-lower histogram), which defined the positive cells (Pos/P2) that internalized the fluorescent probes. The values of MFI presented in this work represent the overall fluorescence of all the single cells (Neg + Pos). Prior detachment of cells for flow cytometry analysis, fluorescent NPs can be visualized by confocal microscopy in cells (C-**D**- HepaRG cells containing fluorescent NPs). In cocultures (**E**) associating GFP expression HepaRG cells (H) and human primary macrophages (M), cells were gated using the SSC-Height versus SSC-Area to identify the single cells (**F**) and the two cell types were segregated using the GFP expression in HepaRG cells detected with the FITC chanel and internalization of DiD-Oil loaded NPs was visualized using the APC chanel (**G**).
